# A modular self-assembling and self-adjuvanting multiepitope peptide nanoparticle vaccine platform to improve the efficacy and immunogenicity of BCG

**DOI:** 10.1101/2024.08.04.606253

**Authors:** Guangzu Zhao, Harindra D. Sathkumara, Socorro Miranda-Hernandez, Julia Seifert, Ana Maria Valencia-Hernandez, Munish Puri, Wenbin Huang, Istvan Toth, Norelle Daly, Mariusz Skwarczynski, Andreas Kupz

## Abstract

After more than a century since its initial development, Bacille Calmette-Guérin (BCG) remains the only licensed vaccine against tuberculosis (TB). Subunit boosters are considered a viable strategy to enhance BCG efficacy, which often wanes in adolescence. While many studies on booster subunit vaccines have concentrated on recombinant proteins, here we developed a novel modular peptide-based subunit vaccine platform that is flexible, cold-chain independent and customizable to diverse circumstances and populations. Each individual peptide building block consists of a linear arrangement comprising a 15-leucine self-assembly inducer moiety, a *Mycobacterium tuberculosis* (Mtb) target epitope and an HLA-E binding moiety, with each moiety separated by a triple lysine spacer. The building blocks, in any combination, were able to form a multiepitope nanoparticle. Six Mtb epitopes were selected to produce the self-assembling and self-adjuvanting peptide-based TB nano-vaccine candidate PNx6. *In vivo* vaccination-challenge experiments demonstrated that subcutaneous boost of parenteral BCG immunization with PNx6 significantly enhanced its immunogenicity and improved its protective efficacy in a murine model of TB by more than 5-fold. Our study present evidence that purely amphiphilic peptides self-assemble into self-adjuvanting nanoparticles with appropriate size and morphology for TB vaccination with great potential for a multitude of other diseases.

## Introduction

Tuberculosis (TB) constitutes a significant global health challenge with an annual incidence of approximately 10 million new cases and 1.5 million fatalities ^1^. Moreover, an estimated one-quarter of the global population harbor latent infections of *Mycobacterium tuberculosis* (Mtb), the causative agent of TB ^2^. The potential reactivation of latent TB infection (LTBI) into destructive pulmonary disease poses a substantial obstacle to any concerted efforts aimed at eradicating TB ^3^. Immunocompromising conditions, notably coinfection with human immunodeficiency virus (HIV) and type 2 diabetes, markedly increase the risk of LTBI reactivation ^4^.

Vaccination remains the cornerstone of infectious disease control by preventing illness and reducing transmission ^5^. So far, Bacille Calmette–Guérin (BCG) developed approximately a century ago, remains the only licensed vaccine for TB ^6^. However, primary neonatal immunization with BCG does not consistently provide protection to adolescents or adults against pulmonary TB ^7^. Furthermore, infant BCG vaccination demonstrated only 37% effectiveness against all forms of TB in children younger than 5 years ^8^. Additionally, the effectiveness of BCG immunization against pulmonary TB varies among adults, and its use is limited in individuals with compromised immunity ^9^. As a live attenuated vaccine, BCG relies on replication and/or persistence to confer immunity. However, exposure to nontuberculous mycobacteria can induce cross-reactive immune responses that hinder BCG replication and diminish its effectiveness ^10^. It is believed that prior exposure could also explain the modest effect of BCG revaccination in previous clinical trials investigating BCG revaccination ^11,12^. Nonetheless, recent studies have demonstrated that the efficacy of BCG revaccination differs across various studies and populations. Some findings suggest modest protection against the development of TB disease, especially in high-risk populations^13^.

In endeavours to address the variability and declining effectiveness of BCG-induced immunity, numerous novel vaccine candidates and modalities are presently under development. In this process, different types of vaccine such as viral vectored vaccines, adjuvant protein subunit vaccines, and whole-cell vaccines incorporating heat-inactivated, fragmented, or genetically modified mycobacteria have been developed ^14–16^. Unlike BCG and other live-attenuated vaccines, non-replicating subunit vaccines based on minimal antigen and synthetic immunostimulatory adjuvants, offer advantages in safety and reduce the adverse effects of vaccination ^17^, which is particularly crucial given the high prevalence of TB in immunocompromised population (i.e., HIV patients) ^18^. To date, together with other types of TB vaccine candidates, only a handful of subunit vaccines have undergone clinical trials and have demonstrated promising efficacy against TB ^15,19–24^. Nonetheless, currently BCG remains the gold-standard and vaccines with higher efficacy are required as outlined in the World Health Organization (WHO) End TB strategy ^25^. Consequently, there has been a growing impetus for the development of new TB vaccine candidates ^26^.

During the activation of an anti-Mtb immune response, macrophages and dendritic cells (DCs) serve as primary antigen-presenting cells (APCs) and are integral for the phagocytosis of Mtb. Upon activation, DCs migrate to lymph nodes where they present mycobacterial antigens on their surface. These antigens are subsequently recognized by classical and donor-unrestricted T cells through major histocompatibility complex (MHC) II and MHC I molecules or CD1-, MR1- or HLA-E-dependent antigen presentation pathways, respectively ^27,28^. Notably, the interaction between T cells that recognise protein-derived antigens and APCs predominantly occurs via peptides rather than entire proteins ^29,30^. Thus, the delivery of immunodominant peptides directly to immune cells via peptide-based vaccines offers a promising approach for vaccine development. The key advantage of peptide-based vaccines lies in their ability to deliver aggregate immunodominant epitopes precisely, thereby minimizing the risk of inducing unwanted immune responses while ensuring the presentation of higher density of antigens compared to other types of vaccine ^31^. Nevertheless, owing to the administration of only the minimal peptide epitopes required for APC to elicit an immune response, peptide-based vaccines often exhibit poor immunogenicity by themselves ^32^. To address this limitation, peptide-based vaccines are commonly administered with adjuvants ^33,34^. Despite the availability of adjuvants for vaccine development, most of them come with inherent drawbacks, including some level of toxicity and the potential to induce adverse effects. These effects may include local reactions at the injection site, such as inflammation, redness, swelling, and pain, as well as systemic reactions like malaise, fever, adjuvant arthritis, and anterior uveitis ^35^. Additionally, the complexity of the preparation process, their often proprietary nature and the associated high costs limit the widespread application of many adjuvants ^36^.

More recently, advancements in nanotechnology have enabled the development of peptide-based vaccines formulated in a nanostructure that have emerged as a promising alternative option ^31^. A previous study demonstrated that a poly-hydrophobic amino acid moiety with a B cell peptide epitope and a T-helper epitope that forms a branch structure is able to self-adjuvant the immunogenicity of the conjugated peptide epitope and induce significantly higher titers of systemic B cell epitope-specific immunoglobulin G (IgG) ^37^. Following analysis involving various amino acids ^38^, it was determined that peptides conjugated with a 15-leucine (15L) poly-amino acid elicited the strongest immune response ^37^. This peptide construct composed of the J8 epitope of group A *Streptococcus* (GAS), the PADRE T-helper epitope and 15L demonstrated the ability to self-assemble into small nanoparticles of 10-30 nm size. These nanoparticles aggregate into bigger chain-like aggregates of nanoparticles (CLANs) of 200-400 nm diameter. Mice vaccinated with this peptide nano-vaccine exhibited higher titers of J8-specific IgG compared to mice immunized with J8 co-administered with complete Freund’s adjuvant (CFA), which is known for its potent adjuvant properties but also associated with high toxicity. Furthermore, mice vaccinated with the peptide nano-vaccine demonstrated nearly complete protection against GAS infection ^37^. Peptide-based strategies have also been employed in the development of vaccines against hookworm ^39^, SARS-CoV-2 ^40^, porcine circovirus 2 ^41^ and cervical cancer ^42^.

Given the difficulties developing effective vaccines for complex pathogens, such as Mtb, it remains critical to diversify the pre-clinical TB vaccine pipeline and to further explore the potential synergy between subunit vaccines and BCG. Protein/adjuvant subunit vaccine development for TB is labour-intensive, slow and requires extensive evaluation of the optimal, often proprietary, adjuvant formulation. Here we developed a novel modular peptide-based subunit vaccine platform that is flexible, cold-chain independent and customizable to diverse circumstances and populations. In this type of vaccine, each basic building block contains a pathogen-derived peptide epitope, a target peptide and a self-assembling and self-adjuvanting inducer peptide. By substituting the pathogen-derived epitope, multiple building blocks can be obtained.

Based on this modular peptide-based subunit vaccine concept, we designed the peptide structure of the basic building block utilizing the 15L motif. Briefly, each individual peptide building block consists of a linear arrangement comprising a 15L moiety, a 15 amino acids peptide TB epitope and a 9 amino acids HLA-E binding moiety, with each moiety separated by a triple lysine spacer. The incorporation of the HLA-E binding peptide aims to allow donor-unrestricted targeting and to broaden the immune cell types that respond to specific Mtb antigens. Six Mtb epitopes were selected to produce 6 individual building blocks. By mixing peptides 1 to 6 we produced the ***p***eptide ***n***anoparticle-based ***6***-epitope TB vaccine candidate PNx6. Here we describe the first application of PNx6 and show that its most effective when administered as a booster to prior BCG vaccination in a murine model of TB.

## Results

### Vaccine design and epitope selection

Based on the self-assembling inducer property of the 15L peptide moiety, a structural blueprint was designed for a single peptide within the framework of the multiepitope peptide-based self-adjuvanting nano-vaccine against Mtb (**Fig. 1A**). Owning to the amphiphilic nature of its configuration, the designed peptide was anticipated to autonomously assemble into nanoparticles within an aqueous milieu. Within this architectural scheme, the 15L peptide moiety and HLA-E binding peptide moiety remain constant, while the epitope moiety undergoes substitution with 6 distinct epitopes, thereby yielding 6 diverse peptides (**Fig. 1B**). The mixture of these 6 peptides will subsequently self-assemble into nanoparticles in phosphate-buffered saline (PBS) (**Fig. 1C**).

**Figure 1.**
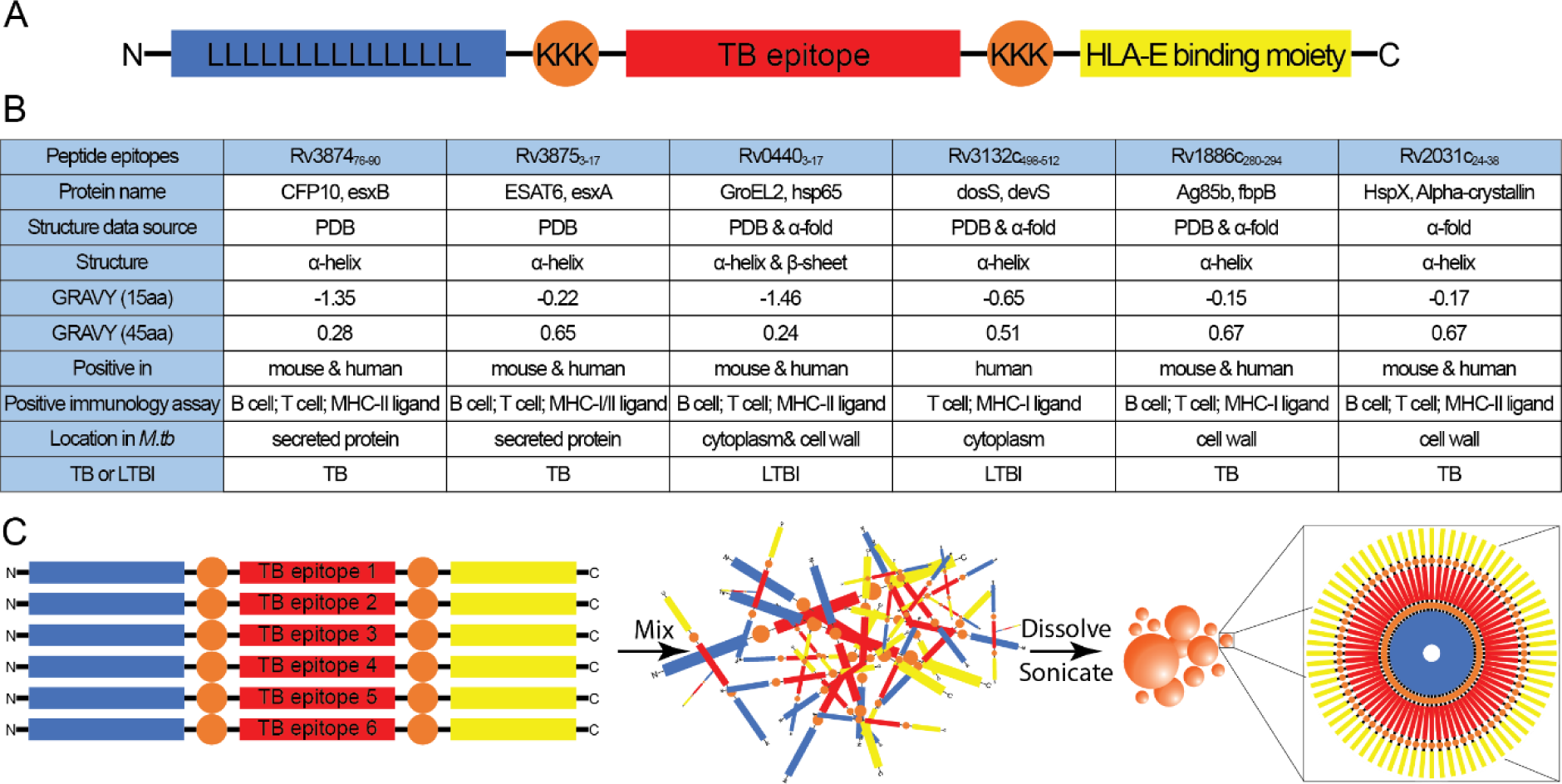
Design of the peptide-based self-assembling nano-vaccine. **A.** Structure design of the individual peptide building blocks. **B.** Details of chosen epitopes **C.** Schematic of peptide-based self-assembling nano-vaccine.

Epitopes were chosen utilizing the Immune Epitope Database (https://www.iedb.org/) according to the following criteria: 1) Epitopes were required to be hydrophilic to render the entire peptide vaccine amphiphilic; 2) The conformation of the peptide epitopes within the original protein should be α-helical, as previous research ^43^ demonstrated that the 15L moiety in the designed peptide vaccine structure could induce the attached peptides to form an α-helix secondary structure; 3) Epitopes were excluded if they contained cysteine residues in their sequences to prevent the formation of disulfide bonds between different peptide molecules; 4) Epitopes should be capable of eliciting an immune response in both mouse models and humans; 5) Epitopes should induce both T cell and B cell responses; 5) The selected epitopes should include epitopes for both active TB and LTBI; 6) The epitopes should be derived from proteins of Mtb that encompass various functions and locations, such as secreted proteins, cell wall proteins, plasma membrane proteins, and cytoplasmic proteins (**Fig. 1B**, **Fig. 2A**).

**Figure 2.**
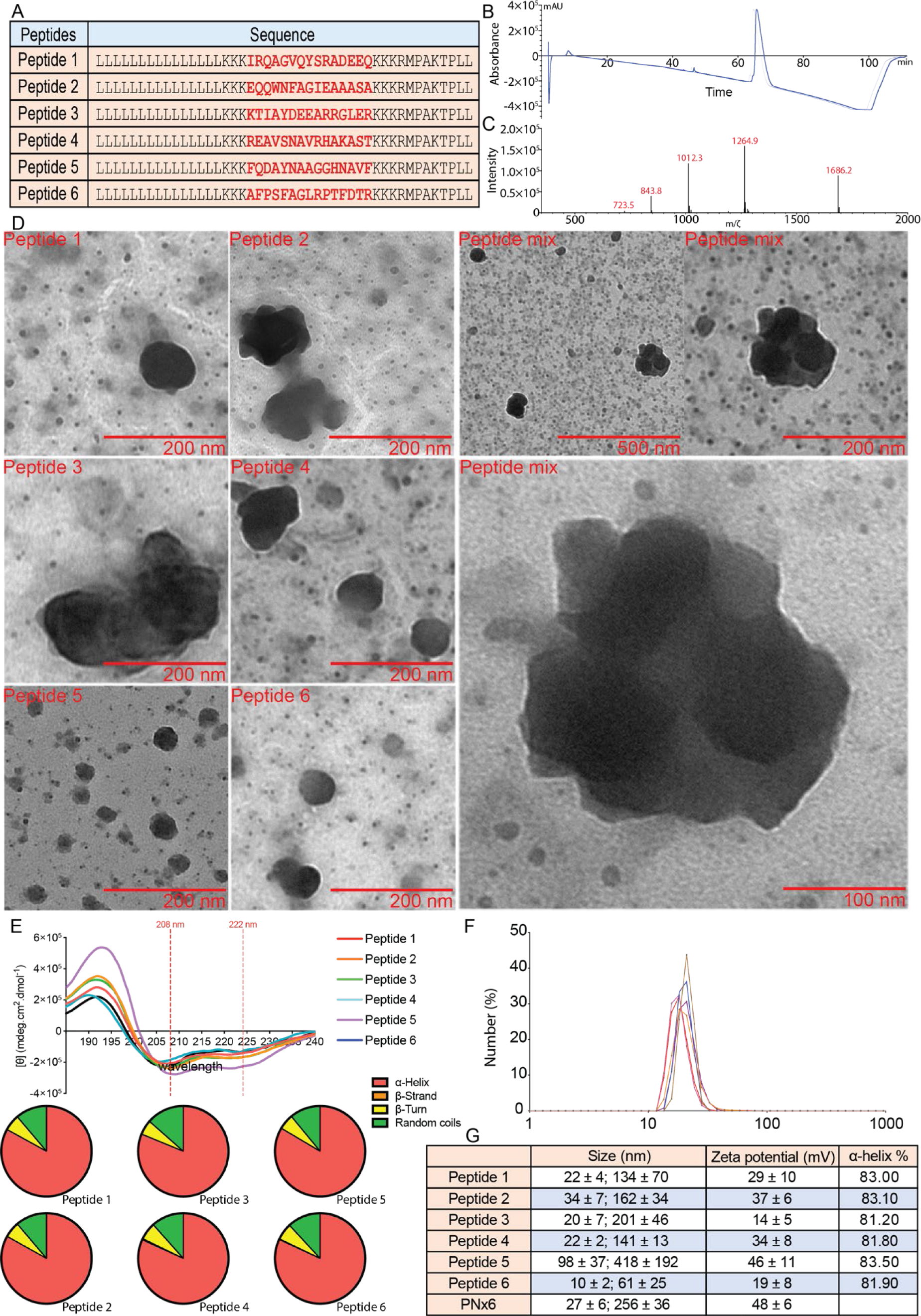
Sequence, purity of synthetic peptides, and size / morphology of self-assembled peptide nanoparticles. **A.** Sequence of peptides for nano-vaccine. Purity of peptides determined by liquid chromatography (LC) and mass spectrometry (MS). **B.** LC and **C.** MS results of epitope 5 as an example of the purity of peptides used in the nano-vaccine formulation. Size and morphology of nanoparticles were determined by TEM. **D.** TEM images showing self-assembled nanoparticles for peptides 1-6 individually and for PNx6, including magnified images of aggregates. **E.** CD results of peptide 1-6. **F.** DLS results of PNx6 by number. **G.** Size, charge and secondary structure details of peptides 1-6 and PNx6.

To inform the sequence of the HLA-E binding epitope, we utilised the findings from a previous study that identified a 9-amino acid (RLPAKAPLL) peptide derived from the Mtb InhA NADH-dependent enoyl reductase protein (Rv1484), which exhibited high-affinity binding with HLA-E proteins not only from mice but also from nonhuman primates (NHP) and humans ^44,45^. However, this 9-amino acid sequence is hydrophobic, while our objective was to incorporate a hydrophilic HLA-E binding peptide into our peptide vaccine building blocks. Fortunately, the study also investigated the binding affinity of the RMPAKAPLL (Leu2Met) and RLPAKTPLL (Ala6Thr) peptides, both of which demonstrated strong binding capacity to the HLA-E protein. Moreover, the RMPAKTPLL (Leu2Met, Ala6Thr) peptide is hydrophilic. Consequently, the RMPAKTPLL peptide was utilized as the HLA-E binding epitope in our project (**Fig. 2A**).

### Synthesised peptides self-assemble into distinct nanoparticles

The synthesis of peptide 1 involved the conjugation of the 15L moiety, the Mtb epitope (Rv3874_76-90_), and the HLA-E binding moiety onto resin utilizing the standard Fmoc-SPPS method. Two triple lysine motifs were incorporated as a spacer and hydrophilicity enhancer between the functional peptide moieties (**Fig. 1A**; **Fig. 2A**). Peptides 2-6 were generated by substituting the Rv3874_76-90_ moiety in peptide 1 with Rv3875_3-17_, Rv0440_3-17_, Rv3132c_498-512_, Rv1866c_280-294_ and Rv2031c_24-38_. All Mtb derived epitopes (Rv3874_76-90_, Rv3875_3-17_, Rv0440_3-17_, Rv3132c_498-512_, Rv1866c_280-294_ and Rv2031c_24-38_) and the HLA-E binding peptide were also synthesized separately through the standard Fmoc-SPPS method to yield epitopes 1-7 that were used for downstream immunological assays. All synthesized peptides have a purity of higher than 95%, as confirmed by LCMS analysis (**Fig. 2B & 2C; Fig. S1A-J; Fig. S2A-N**). During the cleavage of peptides 1-6 from the resin, methionine oxidation occurred. To address this, potassium iodide (KI) (20 equivalents) and ascorbic acid (20 equivalents) were added to the cleavage cocktail, resulting in the reduction of all oxidized methionine residues ^46^. Despite exhibiting very low solubility in water, peptides 1-6 dissolved in PBS, forming a colloid-like solution (**Fig. S3**), with a concentration of >3 mg/mL.

Peptides 1-6 demonstrated the ability to self-assemble into nanoparticles individually in PBS when mixed by vortexing or sonication. Similarly, a mixture of peptides 1-6 (termed PNx6 hereafter) or peptides 1-4 (PNx4; **Fig. S4**) also exhibited the capability to form nanoparticles. The size and morphology of these nanoparticles were assessed utilizing transmission electron microscopy (TEM) (**Fig. 2D**) and dynamic light scattering (DLS) (**Fig. 2F**; **Fig. S5**). Regardless of whether formed individually or as a mixture, the nanoparticles exhibited a similar spherical shape with irregular surface morphology, which is distinct from the CLAN structure which self-assembled by peptides containing a 15L moiety in another study ^37^. Furthermore, the CLAN structure in previous studies are all small nanoparticles (5 – 10 nm) which form 200 - 300 nm aggregates. In contrast, two distinct sizes of nanoparticles formed by individual peptides 1-6 were observed in our study: by number, the majority fell within the range of 10 - 50 nm, while bigger aggregates of 100 - 400 nm were also present (**Fig. 2F&2G**). Various concentrations of peptides in PBS solution were tested, and the morphology and size of the self-assembled nanoparticles remained consistent across the different concentrations. The charge of nanoparticles formed by peptides 1-6 individually and by PNx6 were evaluated by DLS (**Fig. 2G**; **Fig. S6**). All nanoparticles self-assembled by individual peptides had a positive charge of approximately +30 mV except nanoparticles formed by peptides 3 and 6, which charges were ∼+20 mV. PNx6 nanoparticles had the highest charge of ∼+50 mV (**Fig. 2G**). Additionally, the secondary structure of peptides 1-6 were evaluated by circular dichroism (CD) spectroscopy. In line with our expectations based on epitope selection, all peptides formed α-helixes (**Fig. 2E&2G**).

### Subcutaneous boost with PNx6 increases the frequency of CD4^+^ and CD8^+^ T cells

To assess the *in vivo* immunogenicity of PNx6 following parenteral or mucosal vaccination, we used a range of prime-boost vaccination strategies in 6 week-old female C57BL/6 mice. Mice in the positive control group received a single immunization of BCG, administered either subcutaneously (s.c.) or intranasally (i.n.). Mice in the negative control group remained unvaccinated. One group of mice received PNx6 s.c., and one group received PNx6 i.n. at days 0, 14, 28 and 42. In addition, BCG vaccinated mice (both s.c. and i.n.) received an PNx6 boost either s.c. or i.n. at days 14, 28 and 42 (**Fig. 3A&3B**). All 8 vaccination regimens were safe with no adverse reactions observed (data not shown). Mice were sacrificed 60 days post first immunization, and lung, spleen, bronchoalveolar lavage fluid (BALF) and blood were collected for immunological assays (**Fig. 3A**).

**Figure 3.**
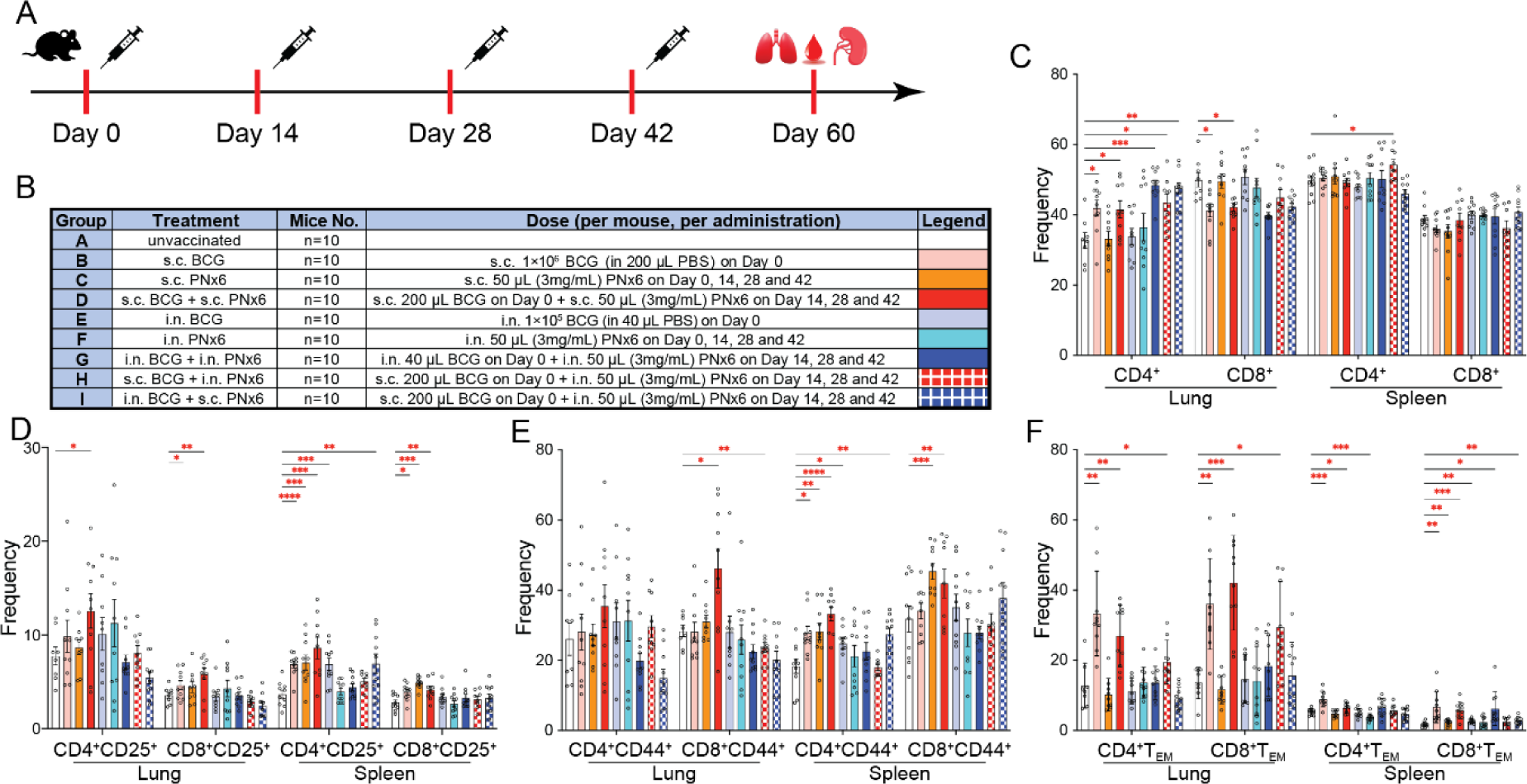
S.c. BCG boosted with s.c. PNx6 increases the frequency of T cells in lung and spleen. **A.**C57BL/6 mice were vaccinated with BCG, PNx6 or BCG boosted with PNx6 either via s.c. or i.n. administration. Unvaccinated mice were used as negative control group. Sixty days later, serum, lung, spleen and BALF were collected. Frequency of immune cells were determined by FACS. **B.** Vaccination details of each group. **C.** Frequency of CD4^+^ and CD8^+^ T cells in lung and spleen of immunized mice. **D.** Frequency of CD4^+^CD25^+^ and CD8^+^CD25^+^ T cells in lung and spleen of immunized mice. **E.** Frequency of CD4^+^ and CD8^+^ memory T cells in lung and spleen of immunized mice. **E.** Frequency of CD4^+^ and CD8^+^ T_EM_ cells in lung and spleen of immunized mice. Each circle in **C**-**F** represents an individual mouse. Differences between groups were tested using the mixed-effects analysis Fisher’s LSD multiple comparison test. All data show means ± SEM or individual mice. *P< 0.05; **P< 0.002; ***P< 0.0002; ****P< 0.0001.

T cell phenotypes in lungs and spleens were assessed by flow cytometry. Immunization with s.c. BCG, s.c. BCG boosted with s.c. PNx6, i.n. BCG boosted with i.n. PNx6, s.c. BCG boosted with i.n. PNx6, and i.n. BCG boosted with s.c. PNx6 led to a notable increase in the frequency of CD4^+^ T cells in the lungs of mice compared to unvaccinated mice. However, the frequency of lung CD8^+^ T cells, as well as both CD4^+^ and CD8^+^ spleen T cells of all vaccinated mice, remained comparable to unvaccinated mice (**Fig. 3C**).

Significantly higher frequencies of CD4^+^ or CD8^+^ CD44^+^ memory T cells in the lungs or spleens of mice vaccinated with s.c. BCG boosted with s.c. PNx6 were observed compared to those only vaccinated with s.c. BCG (**Fig. 3E**). Additionally, the frequency of both CD4^+^ and CD8^+^ effector memory T (T_EM_) cells in the lungs and spleens of mice immunized with either s.c. BCG or s.c. BCG boosted with s.c. PNx6 significantly increased compared to unvaccinated mice (**Fig. 3F**). This trend was also evident in recently activated CD25^+^ CD4^+^and CD8^+^ T cells in lung and spleen (**Fig. 3D**).

### Subcutaneous boost with PNx6 enhances cellular and humoral immunity to BCG

We assessed the induction of IFNγ-secreting cells by PNx6 and BCG boosted with PNx6 using enzyme-linked immunosorbent spot (ELISpot). To this end, splenocytes as well as lung and BALF cells were stimulated with either PNx6 (**Fig. 4A,C&D**) or a mixture of Mtb-derived epitopes 1-6 (**Fig. 4B**). Significantly higher numbers of IFNγ-specific spot-forming cells (SFCs) were detected in the spleen of mice vaccinated with s.c. BCG + s.c. PNx6 boost compared to those vaccinated with s.c. BCG only (**Fig. 4A**). In the lung, the number of IFNγ^+^ T lymphocytes was significantly higher in mice vaccinated with i.n. BCG + i.n. PNx6 boost compared to those receiving i.n. BCG alone (**Fig. 4C**). Similar observations were made in BALF cells (**Fig. 4D**). Although splenocytes from s.c. BCG + s.c. PNx6 vaccinated mice stimulated with epitope 1-6 mixture also produced significantly higher amounts of IFNγ (**Fig. 4B**), the SFCs were still dramatically lower than those stimulated by PNx6 (**Fig. 4A**). Moreover, mice vaccinated with s.c. BCG + i.n. PNx6 exhibited a higher frequency of splenocytes producing IFNγ upon stimulation with PNx6 compared to the s.c. BCG group (**Fig. 4A**). Additionally, mice vaccinated with i.n. BCG + s.c. PNx6 displayed a higher frequency of lung and BALF SFCs upon stimulation with PNx6 compared to the i.n. BCG group (**Fig. 4C&D**). Compared to unstimulated cells, the use of PNx6 resulted in significantly elevated levels of IFNγ^+^ T lymphocytes in the splenocytes of mice receiving s.c. BCG + s.c. PNx6 boost (P < 0.0001), as well as in lung cells (P < 0.002). Additionally, splenocytes were also stimulated with epitopes 1-7 individually (**Fig. S7A-G**). The quantity of SFCs of splenocytes in the s.c. BCG + s.c. PNx6 boost group was also significantly higher than in the unvaccinated group when stimulated by individual epitopes. It appears that epitope 2 (ESAT6_3-17_), epitope 5 (Ag85b_280-294_) and the HLA-E binding epitope are the major stimulants responsible for the increase of IFNγ production while cells stimulated with epitope 1 (CFP10_76-90)_ and 4 (dosS_498-512_) did not lead to a significant increase in SFCs (**Fig. S7A-G, 4E&G**). Moreover, the frequencies of IFNγ^+^ producing cells in lungs from mice vaccinated with s.c. BCG + s.c. PNx6 and BALF from mice immunized with s.c. BCG + i.n. PNx6 upon stimulation with epitope 1-6 mixture were higher than in mice vaccinated with s.c. BCG only (**Fig. S7H-I**).

**Figure 4.**
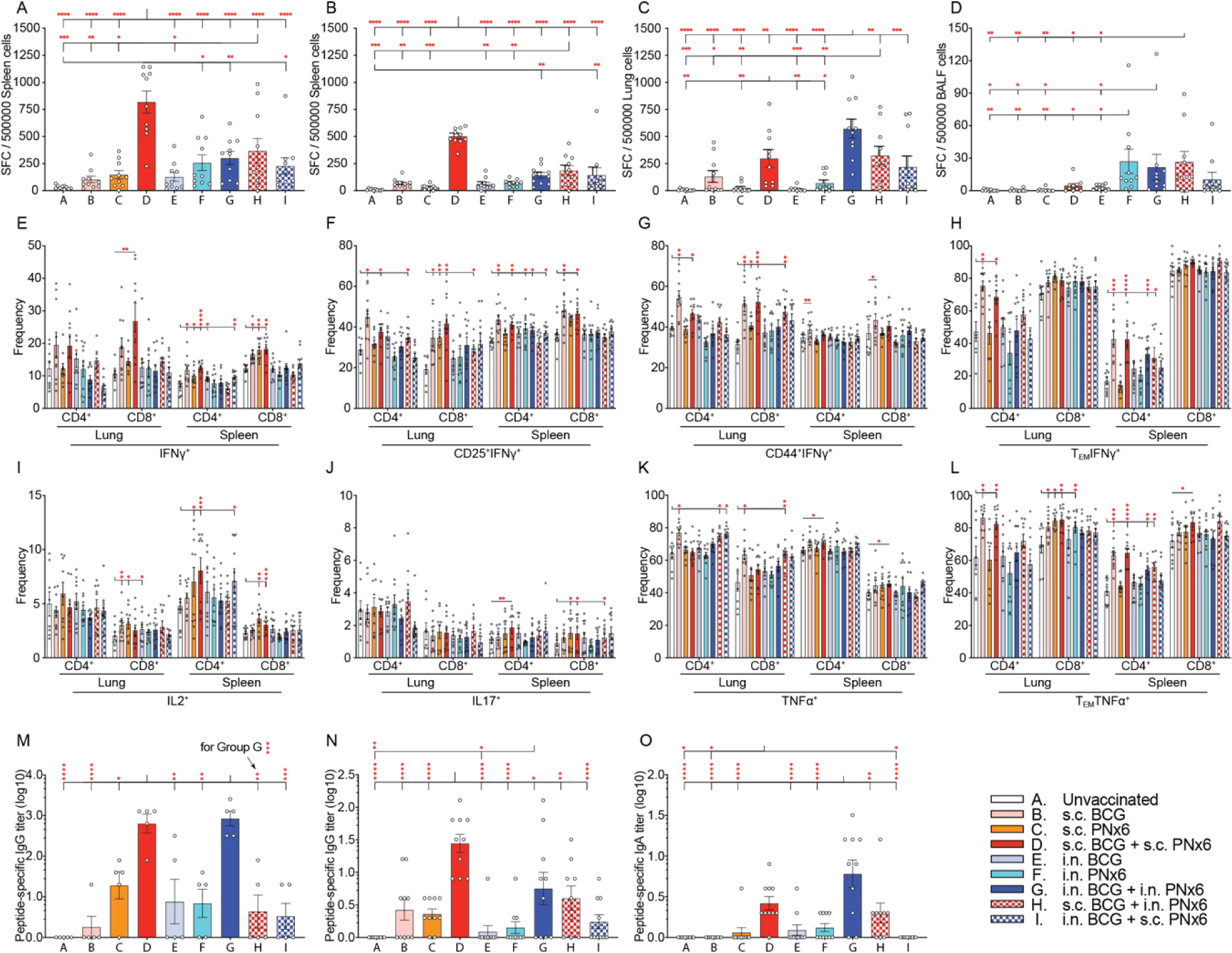
S.c. BCG followed by s.c. PNx6 boost increases cytokine and antibody production. Effect of antigen stimulation on IFNγ secretion was determined by ELISpot. SFC counts (means of two replicates) in IFNγ ELISpot following stimulation of spleen, lung and BALF cells from mice immunized with BCG, PNx6 or BCG boosted with PNx6 by either peptide epitopes or PNx6. Spleen, lung and BALF cells from unvaccinated mice were used as a negative control. **A.** Splenocytes stimulated by PNx6. **B.** Splenocytes stimulated by a mixture of 6 epitopes. **C.** Lung cells stimulated by PNx6. **D.** BALF cells stimulated by PNx6. **E.** Frequency of IFNγ producing CD4^+^ and CD8^+^ lung and spleen T cells. **F.** Frequency of IFNγ producing CD4^+^ and CD8^+^ lung and spleen CD25^+^ T cells. **G.** Frequency of IFNγ producing CD4^+^ and CD8^+^ lung and spleen memory T cells. **H.** Frequency of IFNγ producing CD4^+^ and CD8^+^ lung and spleen T_EM_ cells. **I.** Frequency of IL2 producing CD4^+^ and CD8^+^ lung and spleen T cells. **J.** Frequency of IL17 producing CD4^+^ and CD8^+^ lung and spleen T cells. **K.** Frequency of TNFα producing CD4^+^ and CD8^+^ lung and spleen cells. **L.** Frequency of TNFα producing CD4^+^ and CD8^+^ lung and spleen T_EM_ cells. **M.** Titers of 1-6 epitopes-specific IgG in serum. **N.** Titers of 1-6 epitopes-specific IgG in BALF. **O.** Titers of 1-6 epitopes-specific IgA in BALF. Each circle in **A**-**O** represents an individual mouse. Differences between groups in **A**-**D** were tested using one-way ANOVA with Fisher’s LSD multiple comparison; **E**-**L** were tested using the mixed-effects analysis with Fisher’s LSD test; **M**-**O** were tested using one-way ANOVA with Tukey’s multiple comparison. All data show means ± SEM or individual mice. *P< 0.05; **P< 0.002; ***P< 0.0002; ****P< 0.0001.

In a complimentary approach, intracellular cytokine staining (ICS) was used to assess the cytokine producing capacity of different T cell subsets in lung and spleen (**Fig. 4E-L**; **Fig. S8A-P**). The results revealed that mice vaccinated with s.c. BCG + s.c. PNx6 demonstrated significantly higher frequencies of T cells capable of producing IFNγ compared to unvaccinated mice (**Fig. 4E**). Furthermore, CD8^+^ lung T cells and CD4^+^ and CD8^+^ spleen T cells from this group exhibited higher frequencies of IL-2 production (**Fig. 4I**), while CD4^+^ and CD8^+^ spleen T cells produced higher levels of IL-17 (**Fig. 4J**) and TNFα (**Fig. 4K**). Mice immunized with s.c. BCG or s.c. BCG followed by s.c. PNx6 boost induced significantly higher frequencies of CD4^+^CD25^+^ and CD8^+^CD25^+^ cells producing IFNγ in both lung and spleen (**Fig. 4F**). Additionally, a greater number of memory T cells (both CD4^+^ and CD8^+^) in the lung of mice vaccinated with s.c. BCG or s.c. BCG followed by s.c. PNx6 secreted IFNγ (**Fig. 4G**). Both lung and spleen CD4^+^ T_EM_ cells displayed a higher proportion producing IFNγ when mice were immunized with s.c. BCG or s.c. BCG followed with s.c. PNx6 (**Fig. 4H**). Mice immunized with s.c. BCG followed by s.c. PNx6 boost induced significantly higher amounts of T_EM_ cells (both CD4^+^ and CD8^+^) in both lung and spleen producing TNFα compared to unvaccinated mice (**Fig. 4L**). Mice vaccinated with s.c. BCG exhibited a similar pattern in most T_EM_ cells but displayed no significant difference in spleen CD8^+^ T_EM_ cells compared to the unvaccinated control group (**Fig. 4L**). Additionally, a higher number of both CD4^+^ and CD8^+^ lung memory T cells in mice immunized with s.c. BCG or s.c. BCG followed with s.c. PNx6 were capable of producing TNFα compared to the unvaccinated control group (**Fig. S8N**). Additionally, the levels of almost all measured serum cytokines and chemokines at 60 days post immunization in mice vaccinated with s.c. BCG + s.c. PNx6 were either lower or equivalent when compared to mice immunized with s.c. BCG only. Similarly, circulating serum cytokine and chemokine levels of mice immunized with i.n. BCG + i.n. PNx6 were all lower than mice immunized with i.n. BCG only (**Fig. S9**).

We also assessed serum IgG and BALF IgG & IgA specific to epitopes 1-6 using enzyme-linked immunosorbent assay (ELISA). A noteworthy elevation in epitope 1-6 specific IgG production was observed in all immunized mice. Notably, mice subjected to s.c. vaccination with BCG followed by a s.c. PNx6 boost or i.n. immunization with BCG followed by an i.n. PNx6 boost displayed significantly augmented titers of IgG specific to a mixture of epitopes 1-6 compared to mice administered BCG solely via s.c. or i.n. routes (**Fig. 4M**). Mice vaccinated with s.c. BCG + s.c. PNx6 also had significantly higher titers of epitope 1-6 specific IgG and IgA in BALF when compared to that of unvaccinated and s.c. BCG only mice (**Fig. 4N&O**).

### Subcutaneous boost with PNx6 induces polyfunctional T cells

We also found a notable increase in the frequency of polyfunctional lung CD8^+^ T cells producing quadruple cytokines (IFNγ^+^, IL-2^+^, IL-17^+^ and TNFα^+^) in mice vaccinated with s.c. BCG followed by s.c. PNx6 boost compared to those in the unvaccinated control group (**Fig. 5D&E**). A similar trend was observed in CD4^+^ and CD8^+^ splenocytes (**Fig. 5G, H, J&K**). Concurrently, mice vaccinated with s.c. BCG only exhibited a higher abundance of quadruple cytokine-producing cells in spleen CD8^+^ T cells (**Fig. 5J&K**). Moreover, within the same group, there was a greater proportion of spleen CD4^+^ and CD8^+^ T cells producing triple cytokines in two combinations (IFNγ^+^, IL-2^−^, IL-17^+^ and TNFα^+^; IFNγ^+^, IL-2^+^, IL-17^−^ and TNFα^+^) compared to the negative control group, while mice immunized with s.c. BCG only displayed a higher frequency of CD4^+^ splenic T cells positive for triple cytokines (IFNγ^+^, IL-2^−^, IL-17^+^ and TNFα^+^; **Fig. 5G&H**). Furthermore, mice immunized either with s.c. BCG or s.c. BCG + s.c. PNx6 generated a significantly higher number of double cytokine (IFNγ^+^, IL-2^−^, IL-17^−^ and TNFα^+^) producing CD4^+^ and CD8^+^ T cells in the lung and spleen (**Fig. 5A, B, D, E, G, H, J&K**). Mice immunized with s.c. BCG + s.c. PNx6 also exhibited a significantly higher number of double cytokines in two combinations (IFNγ^−^, IL-2^−^, IL-17^+^ and TNFα^+^; IFNγ^+^, IL-2^−^, IL-17^+^ and TNFα^−^) in both CD4^+^ and CD8^+^ T cells in the spleen (**Fig. 5G, H, J&K**). Additionally, mice immunized with s.c. BCG + s.c. PNx6 also showed a significantly higher frequency of total multifunctional cytokines producing CD4^+^ and CD8^+^ T cells (cells that produce at least two cytokines) in lung and spleen when compared to that of unvaccinated mice (**Fig. 5C, F, I&L**). Moreover, the frequency of total multifunctional CD8^+^ T cells from lungs of mice vaccinated with s.c. BCG + s.c. PNx6 was significantly higher compared to mice immunized with s.c. BCG only (**Fig. 5F**).

**Figure 5.**
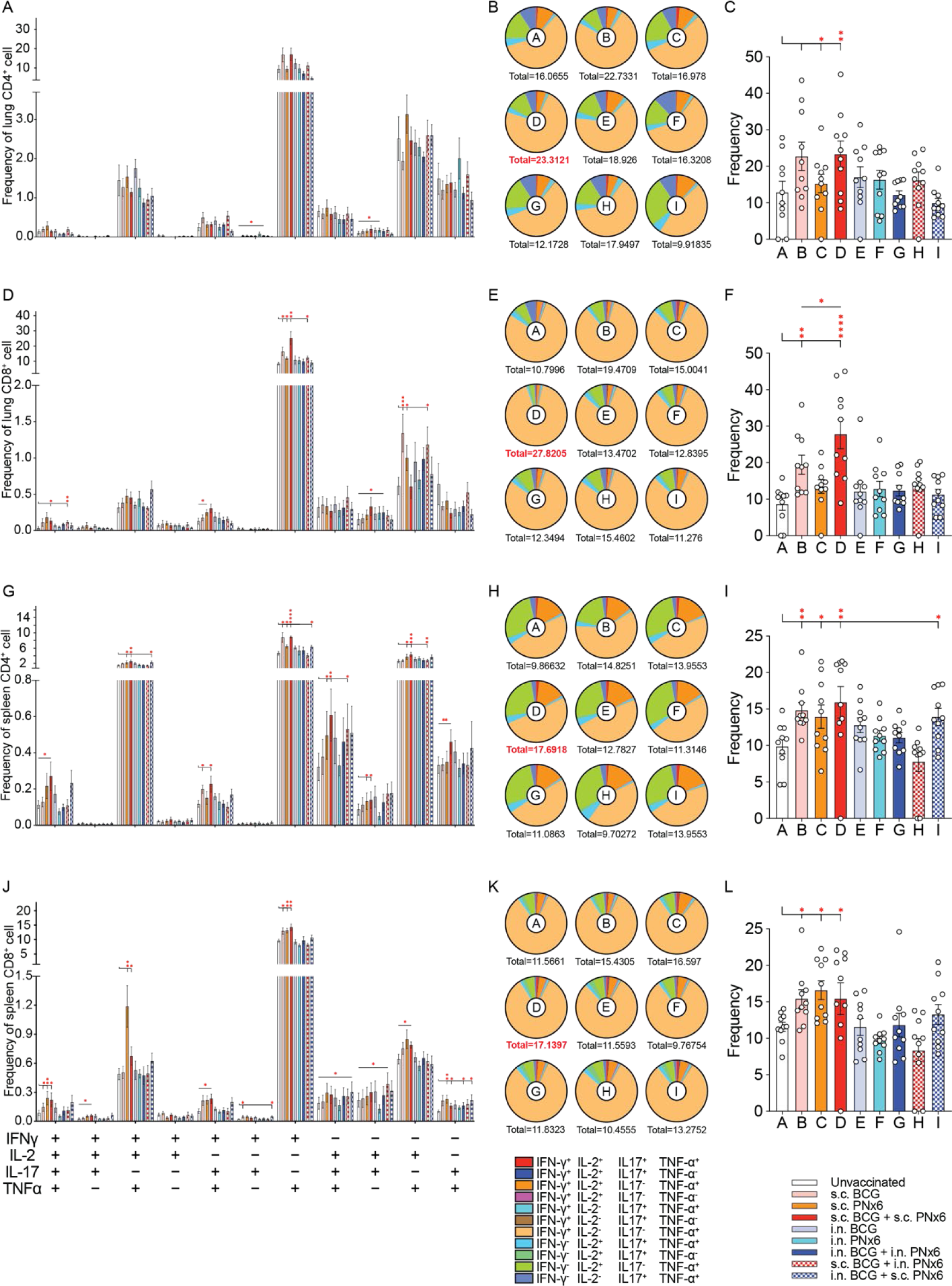
S.c. PNx6 boost of BCG vaccination induces higher frequencies of polyfunctional T cells. **A, D, G, J.** Frequency of polyfunctional cytokine secreting CD4^+^ (A, G) and CD8*^+^* (D, J) T cells in lung and spleen arranged by different cytokine combinations. **B, E, H, K.** Frequency of polyfunctional cytokine secreting CD4^+^ (B, H) and CD8+ (E, K) T cells in lung and spleen arranged by vaccinated groups. **C, F, I, L.** Frequency of total polyfunctional cytokine secreting CD4^+^ (C, I) and CD8+ (F, L) T cells in the lung and spleen. Each circle in **C**, **F**, **I** and **L** represents an individual mouse. Differences between groups in **A**, **D**, **G** and **L** were tested using the mixed-effects analysis with Fisher’s LSD test. **M**-**O** were tested using one-way ANOVA with Fisher’s LSD test. All data show means ± SEM or individual mice. *P< 0.03; **P< 0.002; ***P< 0.0002; ****P< 0.0001.

Collectively these data reveal that a s.c PNx6 boost of prior BCG vaccination not only increases the number and functionality of both CD4^+^ and CD8^+^ T cells, but also leads to a significantly enhanced humoral immune response compared to s.c. BCG vaccination alone.

### Subcutaneous boost with PNx6 provides enhanced protection against Mtb challenge

To assess the protective efficacy of PNx6 against Mtb challenge, C57BL/6 mice were vaccinated with either BCG alone, PNx6 alone, or a combination of BCG and PNx6 boost via s.c. or i.n. routes as described above. At 60 days post initial immunization, all mice were subjected to aerosol challenge with a low dose (50-150 CFU) of Mtb H37Rv. 45 days post-challenge, mice were euthanized and assessed for signs of TB disease (**Fig. 6A**).

**Figure 6.**
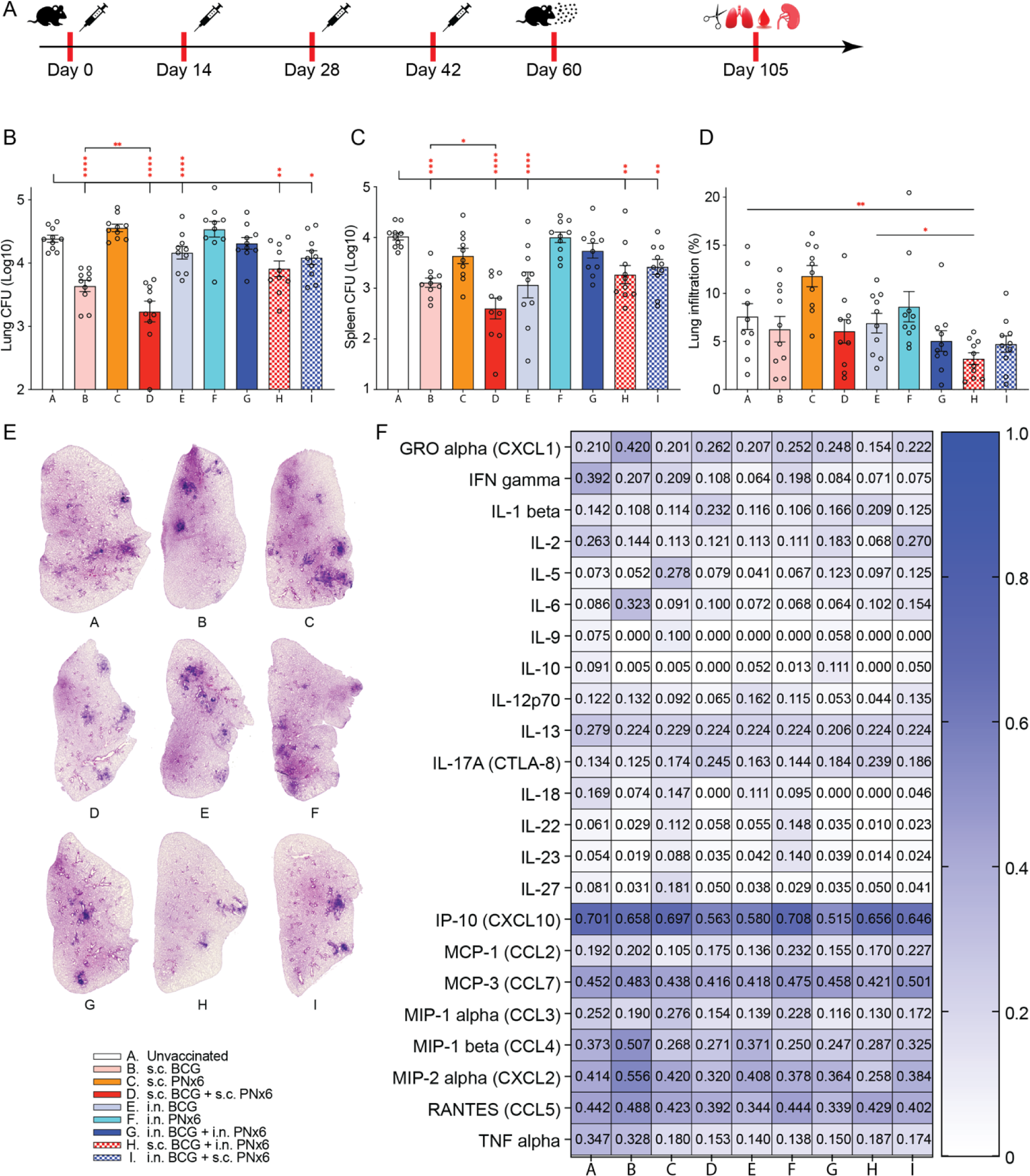
S.c. PNx6 boost of BCG vaccination confers enhanced protection against TB. **A.** C57BL/6 mice were vaccinated with BCG, PNx6 or BCG boosted with PNx6. Sixty days later, mice were aerosol infected with 50-150 CFU of virulent Mtb H37Rv. Mtb bacterial burden was determined 45 days post infection in lungs and spleen. Lung damage was determined as the percentage of the area of dense cell infiltration in H&E-stained lung sections. **B.** Bacterial burden in lung. **C.** Bacterial burden in spleen. **D.** Cell infiltrates in lung. **E.** Representative images of lung infiltration. **F.** Profile of cytokines and chemokines in serum. Each circle in **B**-**D** represents an individual mouse. Differences between groups were tested using the one-way ANOVA with Tukey’s multiple comparison. All data show means ± SEM or individual mice. *P< 0.05; **P< 0.002; ***P< 0.0002; ****P< 0.0001.

Following aerosol Mtb H37Rv challenge, a significant reduction of pathogen burden in both lung and spleen was observed in mice vaccinated with s.c. BCG, i.n. BCG, s.c. BCG + s.c. PNx6, s.c. BCG + i.n. PNx6, and i.n. BCG + s.c. PNx6, compared to unvaccinated mice (**Fig. 6B&C**). Particularly noteworthy, the pulmonary Mtb bacterial burden in mice vaccinated with s.c. BCG + s.c. PNx6 boost was approximately 0.4 log lower (P = 0.0051) than that in mice vaccinated with s.c. BCG only. The CFU count in the spleen of the s.c. BCG + s.c. PNx6 group was also significantly lower (0.5 log; P = 0.02) than that of the s.c. BCG group. Of note, administering PNx6 therapeutically after Mtb infection (**Fig. S10A-C**) or using a PNx4 construct consisting of only 4 peptides (peptide 1-4) as a prime vaccine (**Fig. S10D-F**), did not lead to CFU reduction relative to the s.c. BCG group.

The left lung lobe of each mouse was used for histopathology assessment. The mean tissue infiltration was 7.58% in unvaccinated mice, 6.27% in s.c. BCG vaccinated mice; 11.80% in s.c. PNx6 vaccinated mice; 6.06% in s.c. BCG + s.c. PNx6 vaccinated mice, 6.91% in i.n. BCG vaccinated mice; 8.62% in i.n. PNx6 vaccinated mice; 5.04% in i.n. BCG + i.n. PNx6 vaccinated mice, 3.21% in s.c. BCG + i.n. PNx6 vaccinated mice, 4.73% in i.n. BCG + s.c. PNx6 vaccinated mice, respectively (**Fig. 6D&E**). Although mice vaccinated with s.c. BCG + i.n. PNx6 showed the lowest lung infiltration amongst all groups, the overall pattern of histopathology largely correlated with the pattern of CFU reduction observed in the lung and spleen (**Fig. 6B&C**).

Additionally, at 45 days post-Mtb challenge, mice vaccinated with s.c. BCG followed by an s.c. PNx6 boost demonstrated either comparable or lower serum cytokine and chemokine levels compared to the s.c. BCG-only group. A similar trend in serum cytokine and chemokine levels were observed among mice immunized with i.n. BCG + i.n. PNx6 and i.n. BCG only group (**Fig. 6F**).

Collectively, these results indicate that a s.c. PNx6 boost of BCG vaccination significantly reduces the severity of murine TB in terms of bacterial burden, histopathology, and levels of circulating inflammatory cytokines.

## Discussion

The development of effective vaccines against TB remains a global health priority, particularly in the face of increasing antibiotic resistance and the persistence of TB as a leading cause of morbidity and mortality worldwide. In this study, we explored the potential of a peptide-based multi-epitope nano-vaccine, designed to self-assemble and self-adjuvant, as a novel tool for TB prevention. Our results demonstrate that a parenteral PNx6 boost of prior BCG vaccination enhances anti-TB immunity and leads to significantly improved protective efficacy in a murine model of TB.

It is becoming increasingly clear that an effective TB vaccine will not only need to consist of *multiple* epitopes to induce a broad immune response, but also of the *right* epitopes to elicit a robust and targeted immune response against different stages of Mtb infection. Here we further developed and optimised a peptide-based vaccine platform by incorporating novel components into the structure to enhance immunogenicity. Departing from the T helper epitope utilized in previous peptide nano-vaccine constructs, we instead introduced an HLA-E binding moiety. This HLA-E binding peptide was derived from the Rv1484 Mtb protein, with documented ability to bind to HLA-E molecules across humans, non-human primates, and mice, thus facilitating potential clinical translation ^44^. To generate a linear version of the critical 15-leucine moiety, two triple lysine motifs were included as spacers in our peptide design, as poly-lysine has demonstrated the trend to induce α-helix formation ^43^. Furthermore, the hydrophilicity and positive charges introduced by lysine to the peptide vaccine constructs aimed to improve *in vivo* stability and delivery efficiency, particularly via mucosal administration routes, and to thwart aggregation ^47,48^. Importantly, the selected immunogenic peptide epitopes in our study were predominantly derived from Mtb, rather than BCG, with the aim to minimise the risk of inducing immune responses that control the multiplication of BCG ^49^. Furthermore, adhering to our epitope selection criteria, the combination of peptide epitopes significantly broadens antigenic coverage not only spatially (across different native locations of proteins) and temporally (across TB and LTBI) but also in terms of adaptive immune responses (humoral and cell-mediated; MHC-I & MHC-II) compared to current TB subunit vaccine candidates ^15^. Notably, CFP-10 and ESAT-6 proteins are present as a complex in Mtb secreted proteins, and act as a regulator of macrophage cell death at different stages of Mtb infection ^50^. Hence, we have incorporated epitopes from both proteins to enhance the induced specific immune response against them. Additionally, the peptide epitope derived from the CFP-10 protein harbors a YxxxD/E motif in its sequence, which serves as a secretion signal of ESX-1 substrates found in the PE family of Mtb proteins ^51^. Therefore, the inclusion of this peptide epitope into PNx6 may also induce immune responses against other PE proteins that harbour the YxxxD/E motif. Remarkably, in contrast to other peptide vaccine candidates belonging to the 15L-family that self-assemble into a CLAN structure ^37,43,52^, the PNx6 nanoparticles are closer in similarity to a conventional nanoparticle, characterized by irregular globular shapes. Our observations revealed the presence of two distinct sizes of nanoparticles: the predominant particles ranged between 10-50 nm, with additional larger particles measuring between 100-400 nm. This disparity in morphology may be due to the optimized structure of our peptide design.

One of the notable findings elucidated in this study is the robust adaptive immune response elicited by s.c. BCG followed by s.c. PNx6 boost. Consistent with previous reports on 15L-based nano-vaccine formulations ^37^, we observed that the s.c. BCG + s.c. PNx6 administration resulted in a significant elevation in the titer of epitope specific IgG in serum. Importantly, our investigation confirmed the capacity of the 15L-based nano-vaccine to induce the production of epitope specific IFNγ by spleen, lung and BALF cells. Notably, despite the systemic nature of s.c. BCG + s.c. PNx6 immunization, the production of epitope specific IFNγ in the lung and BALF cells, along with the high titer of IgG and IgA in BALF, suggests that the immune response initiated by parenteral PNx6 delivery may also impact pulmonary sites. This is important, as the induction of pulmonary immune responses via mucosal vaccine delivery has been a focus of TB vaccine research over the last decade ^53^. Our study revealed a higher abundance of CD4^+^ and CD8^+^ memory T cells, particularly T_EM_ cells in the lungs of mice vaccinated with s.c. BCG + s.c. PNx6 compared to unvaccinated mice and compared to s.c. BCG vaccination. Furthermore, CD4^+^ T_EM_ cells in both the spleen and lung from s.c. PNx6 boosted mice exhibited the highest proportion of cells producing both IFNγ and TNFα. It is important to note that i.n. BCG + i.n. PNx6 immunization also elicited a high titer of epitope specific serum IgG and more IFNγ-producing cells in lung and BALF. However, the frequency of CD4^+^ and CD8^+^ memory T cells, and T_EM_ specifically, in mice immunized with i.n. BCG + i.n. PNx6 did not statistically differ from that of unvaccinated mice. Similarly, the production of cytokines such as IFNγ, IL-2, IL-17, and TNFα by memory T cells and T_EM_ cells did not statistically differ compared to unvaccinated mice, suggesting that i.n. BCG + i.n PNx6 provides lower immunity to mice than s.c. BCG + s.c. PNx6. Furthermore, upon stimulation with PNx6, the production of IFNγ by splenocytes of mice immunized with s.c. BCG + i.n. PNx6 was higher than that of mice immunized with either s.c. or i.n. BCG alone, or i.n. BCG + s.c. PNx6, but was lower than that of mice immunized with s.c. BCG + s.c. PNx6, indicating that subcutaneous administration may be the optimal delivery route for PNx6. Given the potential logistical challenges associated with translating safe mucosal vaccine delivery to humans, a parenteral PNx6 boost of existing BCG immunity may be associated with fewer regulatory hurdles and faster implementation.

Mice immunized with s.c. PNx6 produced higher epitope specific antibodies than s.c. BCG administration, but lower levels of IFNγ in T_EM_ cells. As the challenge study confirmed that mice immunized with s.c. PNx6 harboured significantly more Mtb bacteria in both the lung and spleen, our study also suggests that in mice the level of IFNγ-producing T cells provides a more robust immune correlate for evaluating TB vaccine efficacy compared to antibodies. Interestingly, the frequency of total CD8^+^ lung T cells and total CD4^+^ and CD8^+^ spleen T cells did not differ between vaccinated and unvaccinated mice. However, the frequency of IFNγ- and IL-2-producing T cells was significantly higher in mice vaccinated with s.c. BCG + s.c. PNx6 compared to unvaccinated mice. Moreover, the frequency of multifunctional cytokine-producing cells within this group was also significantly higher than that of unvaccinated mice. Overall, mice immunized with s.c. BCG + s.c. PNx6 not only exhibited a stronger cellular and humoral immune response, but also elicited more polyfunctional T cells compared to BCG vaccination alone. These findings support the recent report about synergistic effects of TB vaccine candidate H107 co-administration with BCG ^54^.

While the immunogenicity of a vaccine is important, the protective efficacy against a challenge with the target pathogen remains the critical parameter that determines whether progression to clinical trials is warranted. Our data reveal that the superior immunogenicity of a s.c. PNx6 boost of s.c. BCG vaccination also correlated with the lowest burden of Mtb bacteria in both the lung and spleen. Importantly, the Mtb burden in the lung of mice immunized with s.c. BCG + s.c. PNx6 was significantly lower than that of the s.c. BCG along group. These results align with the reported abilities to reduce Mtb burden in mice of some of the most promising TB vaccine candidates that are currently undergoing clinical trials ^15^. However, the bacterial burden in the lung and spleen of mice vaccinated with i.n. BCG + i.n. PNx6 showed no statistical difference compared to the negative control group, which was unexpected. Furthermore, the efficacy of i.n. BCG + s.c. PNx6 and s.c. BCG + i.n. PNx6 was superior to the placebo group, but not better than s.c. BCG or i.n. BCG alone, suggesting that subcutaneous administration of PNx6 is pivotal for enhancing protective efficacy. These findings are supported by the fact that s.c. BCG + s.c. PNx6 vaccination resulted in one of the lowest lung histopathological infiltration scores and low levels of inflammatory chemokines and cytokines amongst all vaccination groups.

Although safety was not extensively assessed in this study, it is worth noting that PNx6, being composed of peptides, is expected to be non-cytotoxic and no side effects were observed in any of the mice that received the vaccine. Our previous study also showed that 15L was safe, and that no antibody was produced against 15L ^43^. Additionally, the frequency of TNFα-producing cells in the spleen and lung of mice vaccinated with s.c. BCG + s.c. PNx6 was not statistically different from that of the unvaccinated group or the s.c. BCG group. As TNFα is often regarded as an undesirable inflammation indicator ^55^, this result further supports the safety profile of PNx6, and is in line with previous studies on 15L-family peptides, which have indicated that peptides with 15L have no or mild cytotoxicity ^37^.

Given that PNx6 consists of 6 individual peptide building blocks that assemble into nanoparticles individually or in a mixture, it appears very likely that the number of peptide-building blocks incorporated into the vaccine can be increased and easily adjusted. We believe that this feature makes PNx6 unique and differentiates it from other subunit TB vaccine candidates. This plug-in building block approach will allow for customised vaccine formulations that target region-specific dominant Mtb lineages. It will also be possible to include additional peptides that include more potent epitopes and/or those that target additional antigen presenting cells. From a practical perspective, each peptide building block can be produced by an automatic peptide synthesizer. Furthermore, PNx6 has inherent potential as a boost vaccine for neonatal, adolescent and adult BCG vaccination to enhance immunogenicity and protective efficacy by broadening the immune repertoire. In particular, PNx6 induces strong INFy production against ESAT-6, one of the most dominant Mtb antigens that is not included in BCG but is targeted by multiple TB vaccine candidates that are currently being evaluated ^50^. Together with easy storage at room temperature, cold chain-independent transport, and simple preparation requirements, we believe that PNx6 has strong translational potential. Considering that the selected HLA-E binding peptide has stronger affinity to the human HLA-E molecule compared to that of non-human primates and mice, PNx6-mediated boosting of BCG might provide even better immunogenicity and protective efficacy in human clinical trials than what we have observed in mice.

Finally, the nanoparticles used in this study present the first evidence that purely amphiphilic peptides self-assemble into a nanoparticle with appropriate size and morphology for TB vaccine applications. Based on its flexible composition, this peptide-based self-assembling and self-adjuvanting multiepitope vaccine platform could have great potential for a multitude of other diseases.

## Materials and Methods

### Peptide synthesis and purification

Epitope 1 to 7 and peptides 1 to 6 (**Fig. 1**) were synthesized by the manual Fmoc-SPPS method as described elsewhere ^56^. Briefly, the Fmoc group was deprotected twice (1 × 3 min and 1 × 5 min) with 20% piperidine, and double coupling of amino acids (4.2 equivalent) was applied (1× 15 min and 1× 20 min, room temperature). Amino acids were activated by DIPEA (4 equivalent) and HCTU (6.2 equivalent). Peptides were cleaved by trifluoroacetic acid (95%), triisopropyl silane (2.5%) and MilliQ water (2.5%) cocktail. All peptides were purified by RP-HPLC, and their purities were analysed by LCMS. All peptides were lyophilized and kept at −20°C for later use.

Peptide **1** (LLLLLLLLLLLLLLLKKKIRQAGVQYSRADEEQKKKRMPAKTPLL). Yield: 34%. Molecular weight: 5224.6. ESI-MS [M + 3H]^3+^ m/ζ 1742.5 (calc. 1742.5), [M + 4H]^4+^ m/ζ 1307.3 (calc. 1307.1), [M + 5H]^5+^ m/ζ 1046.0 (calc. 1045.9), [M + 6H]^6+^ m/ζ 871.9 (calc. 871.8), [M + 7H]^7+^ m/ζ 747.5 (calc. 747.4). *t*_R_ = 64.32 min (0 to 100% solvent B; C4 column); purity = 98.38%.

Peptide **2** (LLLLLLLLLLLLLLLKKKEQQWNFAGIEAAASAKKKRMPAKTPLL). Yield: 34%. Molecular weight: 5067.4. ESI-MS [M + 3H]^3+^ m/ζ 1689.9 (calc. 1690.1), [M + 4H]^4+^ m/ζ 1267.7 (calc. 1267.8), [M + 5H]^5+^ m/ζ 1014.5 (calc. 1014.5), [M + 6H]^6+^ m/ζ 845.6 (calc. 845.6). *t*_R_ = 55.58 min (0 to 100% solvent B; C4 column); purity = 96.87%.

Peptide **3** (LLLLLLLLLLLLLLLKKKKTIAYDEEARRGLERKKKRMPAKTPLL). Yield: 34%. Molecular weight: 5281.8. ESI-MS [M + 3H]^3+^ m/ζ 1761.4 (calc. 1761.6), [M + 4H]^4+^ m/ζ 1321.4 (calc. 1321.5), [M + 5H]^5+^ m/ζ 1057.3 (calc. 1057.4), [M + 6H]^6+^ m/ζ 881.3 (calc. 881.3). *t*_R_ = 56.84 min (0 to 100% solvent B; C4 column); purity = 99.67%.

Peptide **4** (LLLLLLLLLLLLLLLKKKREAVSNAVRHAKASTKKKRMPAKTPLL). Yield: 34%. Molecular weight: 5071.5. ESI-MS [M + 3H]^3+^ m/ζ 1691.3 (calc. 1691.5), [M + 4H]^4+^ m/ζ 1268.7 (calc. 1268.9), [M + 5H]^5+^ m/ζ 1015.3 (calc. 1015.3), [M + 6H]^6+^ m/ζ 846.3 (calc. 846.3), [M + 7H]^7+^ m/ζ 725.6 (calc. 725.5). *t*_R_ = 58.81 min (0 to 100% solvent B; C4 column); purity = 99.25%.

Peptide **5** (LLLLLLLLLLLLLLLKKKFQDAYNAAGGHNAVFKKKRMPAKTPLL). Yield: 34%. Molecular weight: 5056.4. ESI-MS [M + 3H]^3+^ m/ζ 1686.2 (calc. 1686.5), [M + 4H]^4+^ m/ζ 1265.0 (calc. 1265.1), [M + 5H]^5+^ m/ζ 1012.3 (calc. 1012.3), [M + 6H]^6+^ m/ζ 843.8 (calc. 843.7). *t*_R_ = 66.51 min (0 to 100% solvent B; C4 column); purity = 95.85%.

Peptide **6** (LLLLLLLLLLLLLLLKKKAFPSFAGLRPTFDTRKKKRMPAKTPLL). Yield: 34%. Molecular weight: 5157.6. ESI-MS [M + 3H]^3+^ m/ζ 1720.0 (calc. 1720.2), [M + 4H]^4+^ m/ζ 1290.3 (calc. 1290.4), [M + 5H]^5+^ m/ζ 1032.5 (calc. 1032.5), [M + 6H]^6+^ m/ζ 860.7 (calc. 860.6). *t*_R_ = 63.42 min (0 to 100% solvent B; C4 column); purity = 99.76%.

Epitope **1** (IRQAGVQYSRADEEQ). Yield: 34%. Molecular weight: 1749.8. ESI-MS [M + H]^+^ m/ζ 1750.8 (calc. 1750.8), [M + 2H]^2+^ m/ζ 875.9 (calc. 875.9), [M + 3H]^3+^ m/ζ 584.3 (calc. 584.3). *t*_R_ = 10.73 min (0 to 60% solvent B; C4 column); purity = 98.32%.

Epitope **2** (EQQWNFAGIEAAASA). Yield: 34%. Molecular weight: 1592.7. ESI-MS [M + H]^+^ m/ζ 1593.4 (calc. 1593.7), [M + 2H]^2+^ m/ζ 797.3 (calc. 797.4). *t*_R_ = 33.2 min (0 to 60% solvent B; C4 column); purity = 96.97%.

Epitope **3** (KTIAYDEEARRGLER). Yield: 34%. Molecular weight: 1807.0. ESI-MS [M + H]^+^ m/ζ 1807.9 (calc. 1808.0), [M + 2H]^2+^ m/ζ 904.4 (calc. 904.5), [M + 3H]^3+^ m/ζ 603.3 (calc. 603.3), [M + 4H]^4+^ m/ζ 452.7 (calc. 452.7). *t*_R_ = 17.01 min (0 to 60% solvent B; C4 column); purity = 98.79%.

Epitope **4** (REAVSNAVRHAKAST). Yield: 34%. Molecular weight: 1596.8. ESI-MS [M + 2H]^2+^ m/ζ 799.3 (calc. 799.4), [M + 3H]^3+^ m/ζ 533.3 (calc. 533.3), [M + 4H]^4+^ m/ζ 400.2 (calc. 400.2). *t*_R_ = 10.23 min (0 to 60% solvent B; C4 column); purity = 95.12%.

Epitope **5** (FQDAYNAAGGHNAVF). Yield: 34%. Molecular weight: 1581.6. ESI-MS [M + H]^+^ m/ζ 1582.5 (calc. 1582.6), [M + 2H]^2+^ m/ζ 791.8 (calc. 791.8), [M + 3H]^3+^ m/ζ 528.2 (calc. 528.2). *t*_R_ = 25.24 min (0 to 60% solvent B; C4 column); purity = 97.32%.

Epitope **6** (AFPSFAGLRPTFDTR). Yield: 34%. Molecular weight: 1682.9. ESI-MS [M + H]^+^ m/ζ 1683.6 (calc. 1683.9), [M + 2H]^2+^ m/ζ 842.4 (calc. 842.5), [M + 3H]^3+^ m/ζ 562.0 (calc. 562.0). *t*_R_ = 31.38 min (0 to 60% solvent B; C4 column); purity = 99.46%.

Epitope **7** (RMPAKTPLL). Yield: 34%. Molecular weight: 1026.3. ESI-MS [M + H]^+^ m/ζ 1026.7 (calc. 1027.3), [M + 2H]^2+^ m/ζ 514.2 (calc. 514.2), [M + 3H]^3+^ m/ζ 343.1 (calc. 343.1). *t*_R_ = 13.68 min (0 to 60% solvent B; C4 column); purity = 98.72%.

### Nanoparticle size measurement

The morphologies of nanoparticles formed by self-assembled peptides were visualized by TEM (HT7800, HITACHI Ltd., JEOL Ltd., Japan) operating at a voltage of 80 kV, and DLS. For TEM analysis, peptide 1-6 individually and PNx6 PBS solution were added to copper 200 mesh grids (Ted Pella) with glow-discharged carbon-coated pretreated and then using 2% phosphotungstic acid (PTA) for negative staining for 1 minute. The remaining staining solution was removed by filter paper attached to the edge of the grid, then the grid was air-dried for 10 minutes before TEM analysis. For the DLS assay, the average particle size (nm) of peptide 1-6 individually and PNx6 in water solution (0.1 mg/mL) were measured by a Nanosizer (Zetasizer Nano Series ZS, Malvern Instruments, United Kingdom) at 25 °C with disposable capillary cuvettes.

### Nanoparticle charge measurement

The average particle charges (mV) of peptide 1-6 individually and PNx6 in water solution (0.1 mg/mL) were measured by a Nanosizer (Zetasizer Nano Series ZS, Malvern Instruments, United Kingdom) at 25 °C with disposable capillary cuvettes.

### CD spectroscopy

Peptides 1-6 were measured using a 1 mm quartz cuvette with 1 mm path length (Starna) at 23 °C on a Jasco J710 spectropolarimeter (JASCO, Japan) and data was collected between 185 nm and 275 nm with a continuous scanning speed of 50 nm/min. The equation of molar ellipticity [θ] = mdeg × 10^3^ / [L(mm) x C(mM) x n] (deg.cm^2^.dmol^−1^) [L: cell path length;(mm); C (peptide concentration) = number of mol/volume (mM); n: number of amino acids] was used to calculate the measured data. The Online CD Analysis platform was employed to conduct a quantitative analysis of the secondary structure content ^57^.

### Mice

Female C57BL/6 mice were purchased from the Animal Resource Centre (ARC; Perth, Australia) and maintained in the biosafety level 2 and 3 animal facilities of the Australian Institute of Tropical Health and Medicine at James Cook University, Australia. All mice were allowed to acclimatize to the new environment for 7 days before any experiments were performed. All experiments were conducted in accordance with the National Health and Medical Research Council (NHMRC) animal care guidelines and were approved by the animal ethics committee (A2837) of James Cook University, Australia.

### Vaccinations

Prior to vaccination, frozen vials of vaccine stocks were thawed, washed and diluted in PBS. Mice were 6 weeks old when they were vaccinated. 9 groups of mice (10 mice each group) were immunized with BCG, PNx6 or BCG boosted with PNx6 either subcutaneously or intranasally. Mice in groups B, D and H were immunised with BCG subcutaneously (approximately 10^6^ CFU of bacteria in 200 µl of sterile PBS) on day 1; mice in group E, G and I were vaccinated with BCG intranasally (approximately 10^5^ CFU of bacteria in 40 µl of sterile PBS) on day 1; mice in group C were administrated with peptide vaccine subcutaneously (300 μg peptides in 40 µl of sterile PBS) on day 1, 14, 28 and 42; mice in group F were immunised with peptide vaccine intranasally (300 μg peptides in 40 µl of sterile PBS) on day 1, 14, 28 and 42; mice in group D and I were boosted with peptide vaccine subcutaneously (300 μg peptides in 40 µl of sterile PBS) on day 14, 28 and 42; mice in group G and H were boosted with peptide vaccine intranasally (300 μg peptides in 40 µl of sterile PBS) on day 14, 28 and 42; mice in group A remained unvaccinated as negative control group.

In a pilot vaccination study, PNx4 was administered either s.c. or i.n. at days 0, 14, 28 and 42 and compared to s.c BCG given once at day 0.

### Cell and serum preparation for ICS, ELISpot and ELISA

On day 60 mice were sacrificed by carbon dioxide asphyxiation and spleen, lung, blood and bronchoalveolar lavage fluid (BALF) were collected.

Lungs were collected in GentleMACS tubes and homognized using a GentleMACS™ Dissociator (Miltenyi Biotec). The cell suspensions were incubated with 7.5 µg/ml Collagenase D (Sigma) and 1.75 µg/ml Collagenase VIII (Sigma) for 30 mins at 37°C and subsequently passed through 70 µm cell strainers. Following incubation with erythrocyte lysis buffer, cells were washed with FACS buffer to obtain single a cell suspension. The cell concentration was counted using a hemocytometer. 0.5×10^6^ cells were prepared using complete media for ICS and ELISpot.

Spleens were mechanically dissociated by passing through 70 µm cell strainers. Cell suspensions were then incubated with erythrocyte lysis buffer and washed with FACS buffer to obtain single cell suspension. Cell concentrations were determined using a CASY cell counter. 0.5×10^6^ cells were prepared using complete media for ICS and ELISpot.

BALF was centrifuged first, then incubated with erythrocyte lysis buffer and washed with FACS buffer. The cell concentrations were counted using a hemocytometer. 0.5×10^5^ cells were prepared using complete media for ELISpot.

Blood was collected into Z-gel tubes (Sarstedt) and serum was prepared by centrifugation of clotted blood at 8000 rpm for 8 minutes and stored at −20°C for ELISA.

### Intracellular cytokine staining (ICS)

Splenocytes or lung cells were cultured in presence or absence of 2 μL/mL of cell stimulation cocktail (eBioscience, 00-4975-93) and brefeldin A (BD, 555029) in RPMI complete media for 6 hours at 37°C in a humidified chamber and 5% CO_2_. The cell surface was stained with the following antibodies and fluorochromes: CD3e (AF700; clone 500A2), CD8e (BV510; clone 53-6.7), CD4 (BUV395; clone GK1.5), CD25 (BV711; clone PC61), CD44 (BV421; clone IM7), CD69 (PE/CF594; clone H1, 2F3), CD62L (PE/Cy7; clone MEL-14), CD103 (APC; clone M290), KLRG (FITC; clone 2F1) first, and then fixed and permeabilized using eBioscience Intracellular Fixation and Permeabilization kit (Invitrogen, 88-8824-00). Cell suspensions were then stained with IFNγ (PerCP/Cy5.5; clone XMG1.2), IL-2 (BV605; clone JES6-5H4) IL-17 (BV786; cloneTC11-18H10) and TNF-α (PE; clone MPG-XT22). Unstained cells, and cells stained with the single antibodies were used as controls for the compensation setup. Stained cells were acquired on a BD Fortessa X20 flow cytometer; 200,000 cells from spleen and lungs were recorded, respectively. Effector memory T (T_EM_) cells were considered as CD44^+^KLRG^+^CD69^−^, central memory T (T_CM_) cells as CD44^+^KLRG^−^ CD69^−^CD103^−^ and resident memory T (T_RM_) cells as CD44^+^CD69^+^CD103^+^ cells.

### IFNγ ELISpot

0.5×10^6^ splenocytes, 0.5×10^6^ lung cells or 0.5×10^5^ BALF cells in 100 µl RPMI complete media were added into each well of purified rat anti-mouse IFNγ (Mabtech, 3321-3-1000) pre-coated ELISpot plates, followed with 50 µl stimulant in RPMI complete media and 50 µl complete media. ELISpot plates were incubated in a humidified chamber and 5% CO_2_ incubator for 42 hours at 37°C. After the incubation, plates were washed with PBST (0.05% tween in PBS v/v), followed with adding 50 µl per well of 1 µg/ml biotin rat anti-mouse IFNγ (Mabtech, 3321-6-1000) into each well and incubated for 2 hours at room temperature. Then plates were washed with PBST and 50 µl of 1 µg/ml streptavidin (Mabtech, 3310-9-1000) was added to each well of ELISpot plates. After 1 hour incubation at room temperature in the dark all plates were washed with PBST and stained with 50 µl TMB (Mabtech, 3651-10). After the spots developed, plates were washed again with PBST and dried overnight. All plates were read using an automated ELISpot reader (Zeiss KS ELISpot).

### Serum IgG ELISA

Epitope specific IgG was analyzed by ELISA. Costar 96 well plates (Corning, USA, 3590) were coated with the 6 peptide epitopes mixture (pH 9.6, 120 µg/ml in a carbonate coating buffer) and incubated at room temperature for 8 h, then washed with PBST and blocked with 150 µl of 5% skim milk PBST. After overnight incubation at 4°C, plates were washed 4 times with PBST and filled with 200 µl 20 times diluted serum followed by serial dilution down the plate in 0.5% skim milk PBST buffer. After 2 hours incubation in a humidified chamber and 5% CO_2_ incubator at 37°C, plates were washed 4 times with PBST, followed with 100 µl per well of either 1:3000 diluted biotin goat anti-mouse IgG (in 0.5% skim milk PBST; Sigma SAB4600004-250µl) or 1:1000 diluted biotin goat anti-mouse IgA (BD, 556978), and then incubated 2 hours at 37°C in the incubator. After incubation, plates were washed 4 times with PBST and filled each well with 100 µl 1000 times diluted HRP-conjugated streptavidin (BD, 554066) in 0.5% skim milk PBST buffer, then incubated for 1.5 hours at 37°C. Plates were washed 4 times after incubation, then 100 µl TMB (BD, 555214) was added into each well and plates were kept in the dark at room temperature for 20 minutes. Reaction was stopped by adding 50 µl of 0.18 M H_2_SO_4_ into each well, and then the absorbance of each well were measured at λ 450 nm by POLARstar Omega plate reader (BMG Labtech).

### Mtb challenge

6 weeks old female mice were vaccinated and boosted as outlined above. 10 groups of mice (10 mice each group) were immunized with BCG, PNx6 or BCG boosted with PNx6 either subcutaneously or intranasally. One group of mice remained unvaccinated as negative control group.

For therapeutic use of PNx6, Mtb-infected mice that had previously been vaccinated with PNx6 either s.c. or i.n., received an additional dose of s.c. or i.n. PNx6 15 days post infection.

The Mtb challenge was conducted within a Biosafety Level 3 (BSL3) laboratory precisely 60 days post-vaccination. Initially, frozen aliquots of Mtb were thawed, subsequently diluted to achieve the desired concentration, and subjected to ultrasound treatment in a water bath to disrupt bacterial aggregates. Mice were then exposed to a dose (50-150 CFU) of Mtb H37Rv utilizing a Glas-Col inhalation exposure system. The initial infectious dose was determined through the plating of lung tissues obtained from five mice one day following aerosol infection onto 10% OADC enriched 7H11 agar plates.

### Serum Cytokine and Chemokine Analysis

Frozen serum samples underwent thawing and preparation in accordance with the specifications outlined in the Bio-Plex Pro Mouse Cytokine 23-Plex assay (BioRad). Subsequently, analysis was performed utilizing a MagPix instrument (Luminex). The data analysis and visualization of cytokine levels were in heatmap format (log_2_ concentration).

### CFU Enumeration

Lung and spleen tissues collected from mice were homogenized using GentleMACS tubes containing 1 mL of sterile PBS/0.05% Tween 80. Serial dilutions of the homogenates were then plated onto 10% OADC-enriched 7H11 agar plates supplemented with 10 μg/mL cycloheximide and 20 μg/mL ampicillin. These agar plates were subsequently sealed and aerobically incubated for a duration of 5 to 6 weeks at 37°C. Following incubation, colonies were enumerated, and the total CFUs per organ were calculated based on the dilution factors and the size of the respective organs.

### Histopathology

The left lung lobes obtained from Mtb infected mice were fixed overnight in a 10% w/v formalin solution, subsequently transferred to 70% ethanol, and then embedded in paraffin. Following this procedure, the processed lung samples were sectioned into 4 µm slices, which were then placed onto glass slides, dewaxed, and stained with hematoxylin and eosin (H&E). These stained slides were subjected to scanning using an Aperio CS2 scanner (Leica), and the resulting images were analyzed utilizing both Image Scope (Leica) and ImageJ software (NIH). To determine the extent of lung damage, the total surface area was measured, followed by the quantification of areas displaying dense cell infiltration, in accordance with previously established protocols.

### Statistics

Statistical analysis was conducted, and graphical representations were produced using Prism version 10.2.1 (GraphPad). The data underwent analysis through either one-way ANOVA accompanied by Tukey’s multiple comparison post-test or mixed-effects analysis with Fisher’s LSD multiple comparison test. A significance threshold of p<0.05 was adopted, unless otherwise specified.

## Author contribution

Conceptualization: AK, GZ. Methodology: AK, GZ. Investigation: GZ, HDS, SMH, JS, AMVH, MP, WH, MS, AK. Intellectual support: ND, IT. Manuscript writing: GZ, AK. Funding acquisition: AK.

## Acknowledgements

We thank Chris Wright, Lachlan Pomfrett, Helma Antony and Serrin Rowarth for assistance with BSL3 and animal facility operations. We thank David Pattinson and Daniel Browne for assistance with ELISpot training, and Jennifer Whan and Jordi Nelis for assistance with TEM. This work was supported by the National Health and Medical Research Council through a NHMRC Ideas (APP2001262) and Investigator Grant (APP2008715) to A.K. M.S was supported by ARC Discovery Project DP21010280.

## Supplementary Material

**Figure S1.**
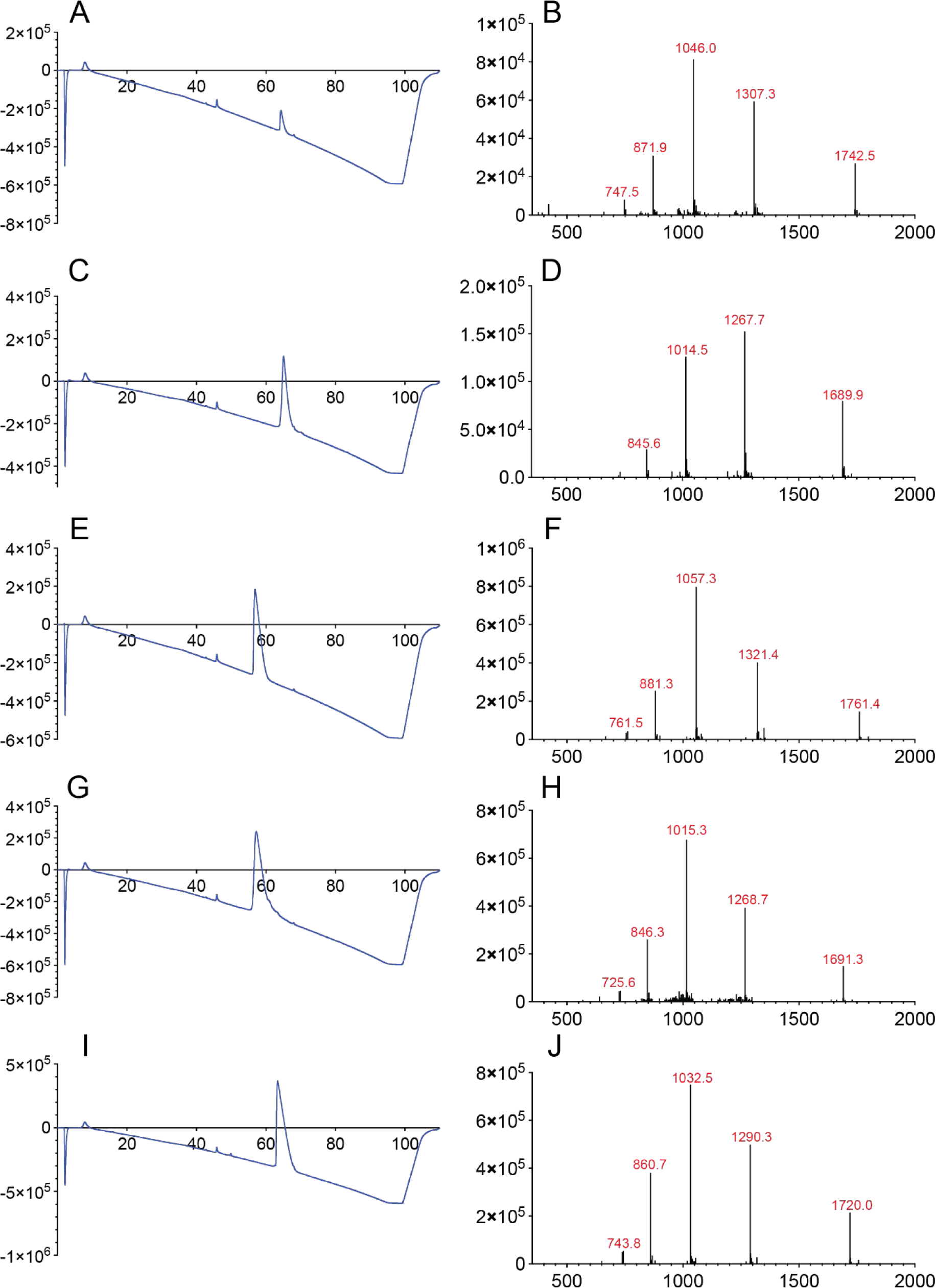
Mass spectrum (MS) and Liquid chromatography (LC) spectra of peptides 1-4 & 6. **A.** LC of peptide 1; **B.** MS of peptide 1; **C.** LC of peptide 2; **D.** MS of peptide 2; **E.** LC of peptide 3; **F.** MS of peptide 3; **G.** LC of peptide 4; **H.** MS of peptide 4; **I.** LC of peptide 6; **J.** MS of peptide 6.

**Figure S2.**
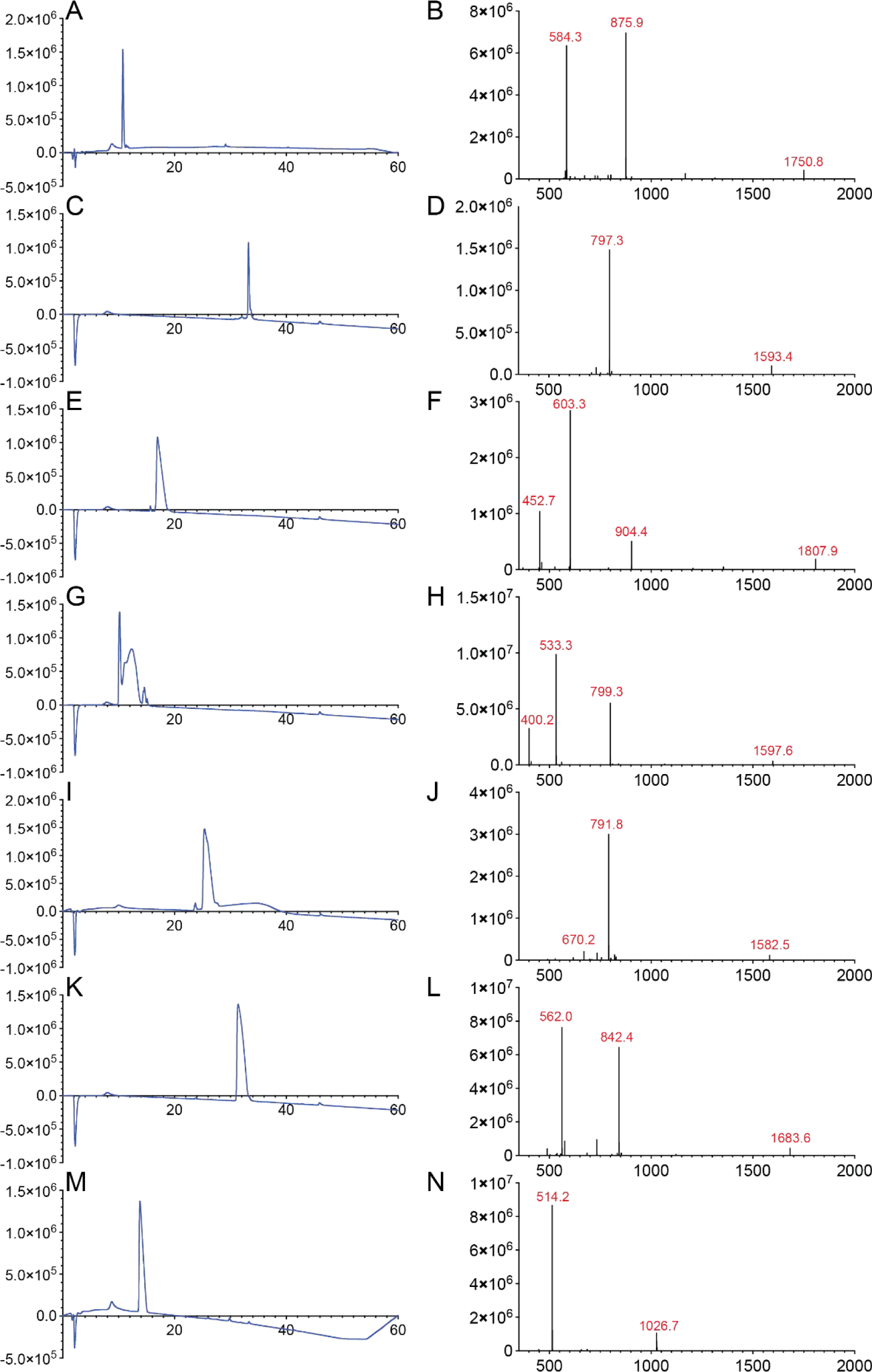
MS and LC spectra of epitopes 1-6 & HLA-E binding peptide. **A.** LC of epitope 1; **B.** MS of epitope 1; **C.** LC of epitope 2; **D.** MS of epitope 2; **E.** LC of epitope 3; **F.** MS of epitope 3; **G.** LC of epitope 4; **H.** MS of epitope 4; **I.** LC of epitope 5; **J.** MS of epitope 5; **K.** LC of epitope 6; **L.** MS of epitope 6; **M.** LC of HLA-E binding peptide; **N.** MS of HLA-E binding peptide.

**Figure S3.**
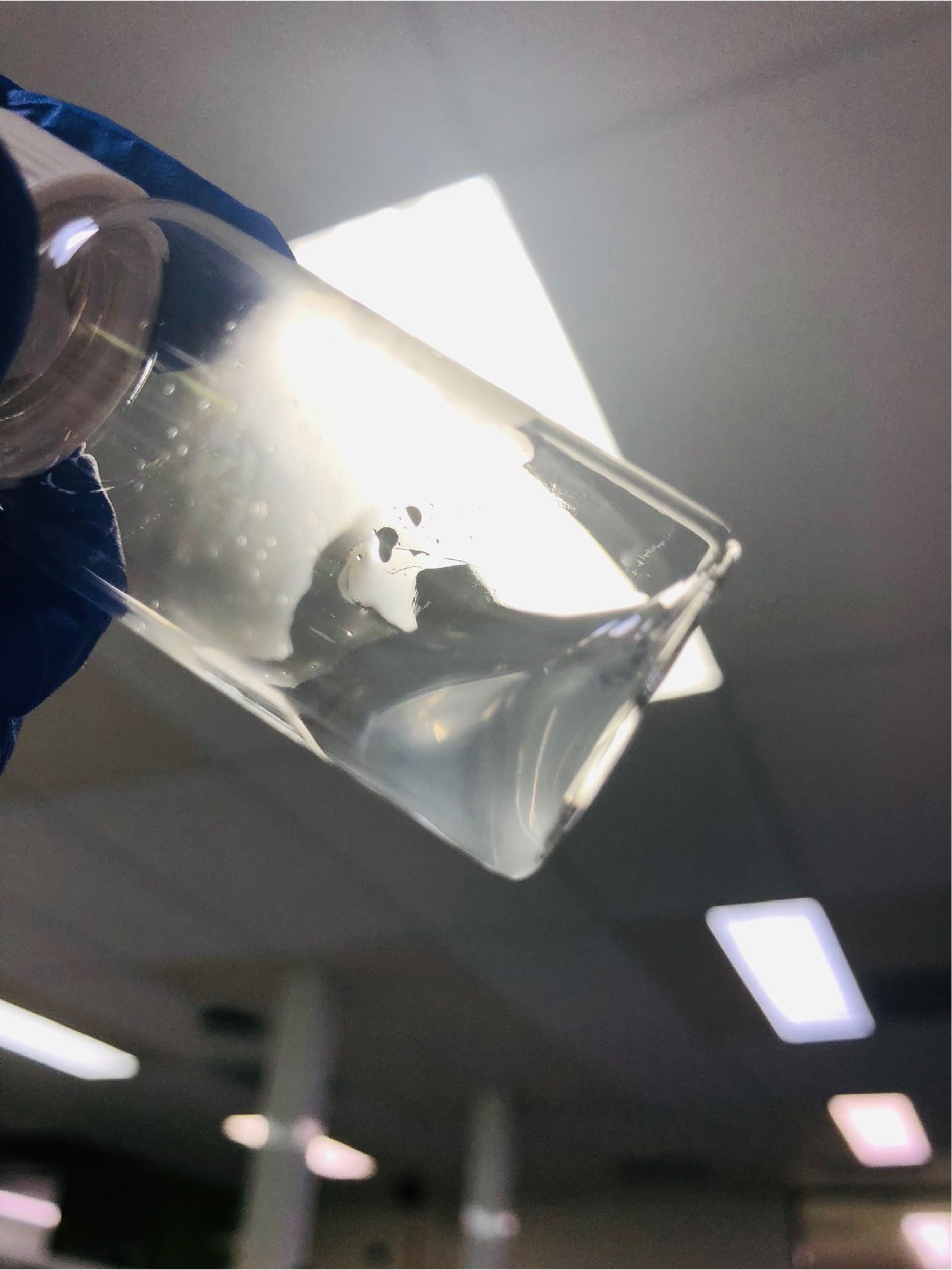
A colloid-like solution formed by the peptide vaccine.

**Figure S4.**
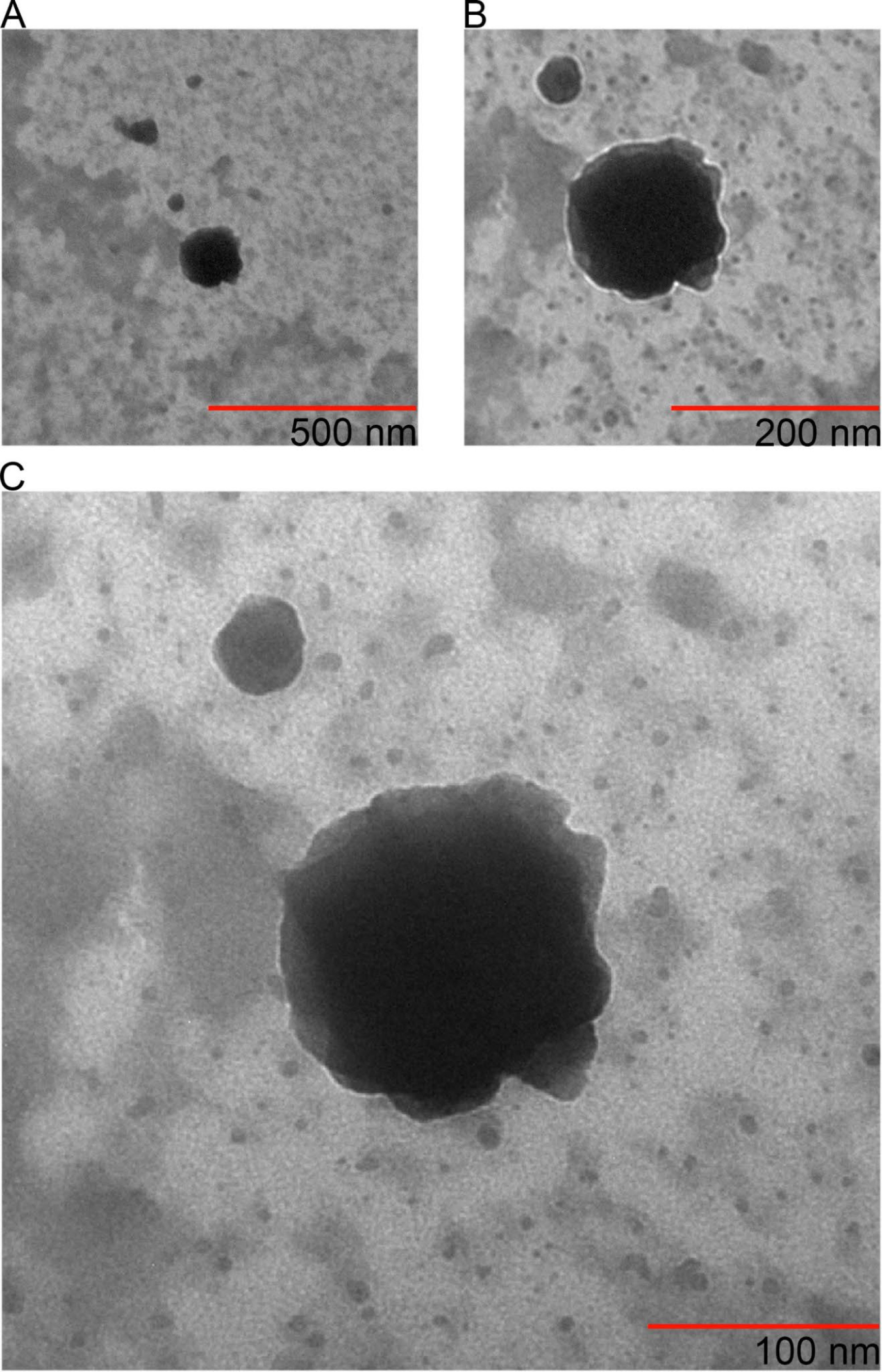
TEM photographs of nanoparticles formed by peptide 1-4 mixture. **A.** Scale bar 500 nm. **B.** Scale bar 200 nm. **C.** Scale bar 100 nm.

**Figure S5.**
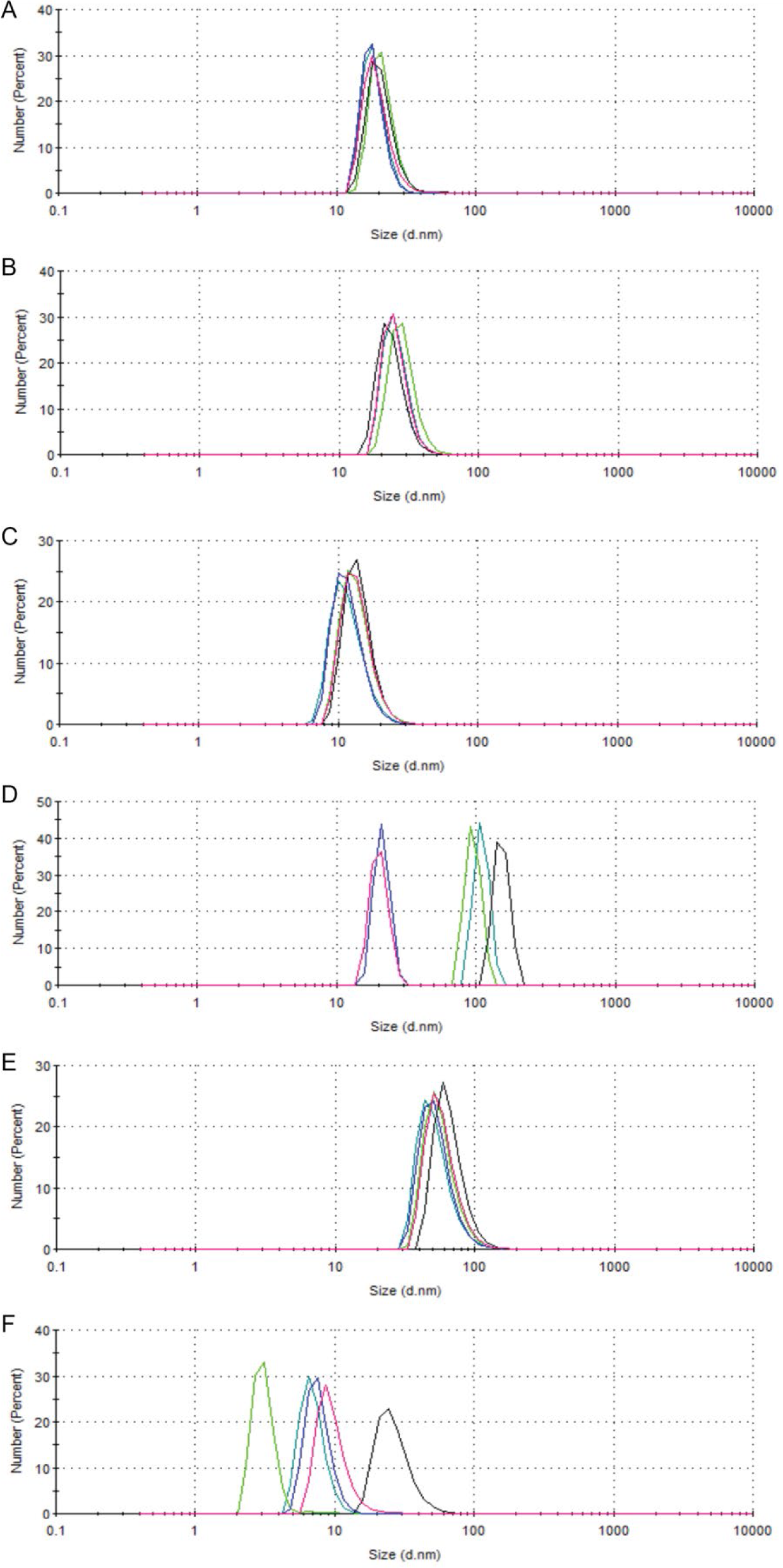
Particle size distribution for nanoparticles self-assembled by peptides 1-6. **A.** particle size distribution of nanoparticles self-assembled by peptide 1 analyzed by number; **B.** particle size distribution of nanoparticles self-assembled by peptide 2 analyzed by number; **C.** particle size distribution of nanoparticles self-assembled by peptide 3 analyzed by number; **D.** particle size distribution of nanoparticles self-assembled by peptide 4 analyzed by number; **E.** particle size distribution of nanoparticles self-assembled by peptide 5 analyzed by number; **F.** particle size distribution of nanoparticles self-assembled by peptide 6 analyzed by number. Five independent replicated measurements per compound recorded by dynamic light scattering (DLS).

**Figure S6.**
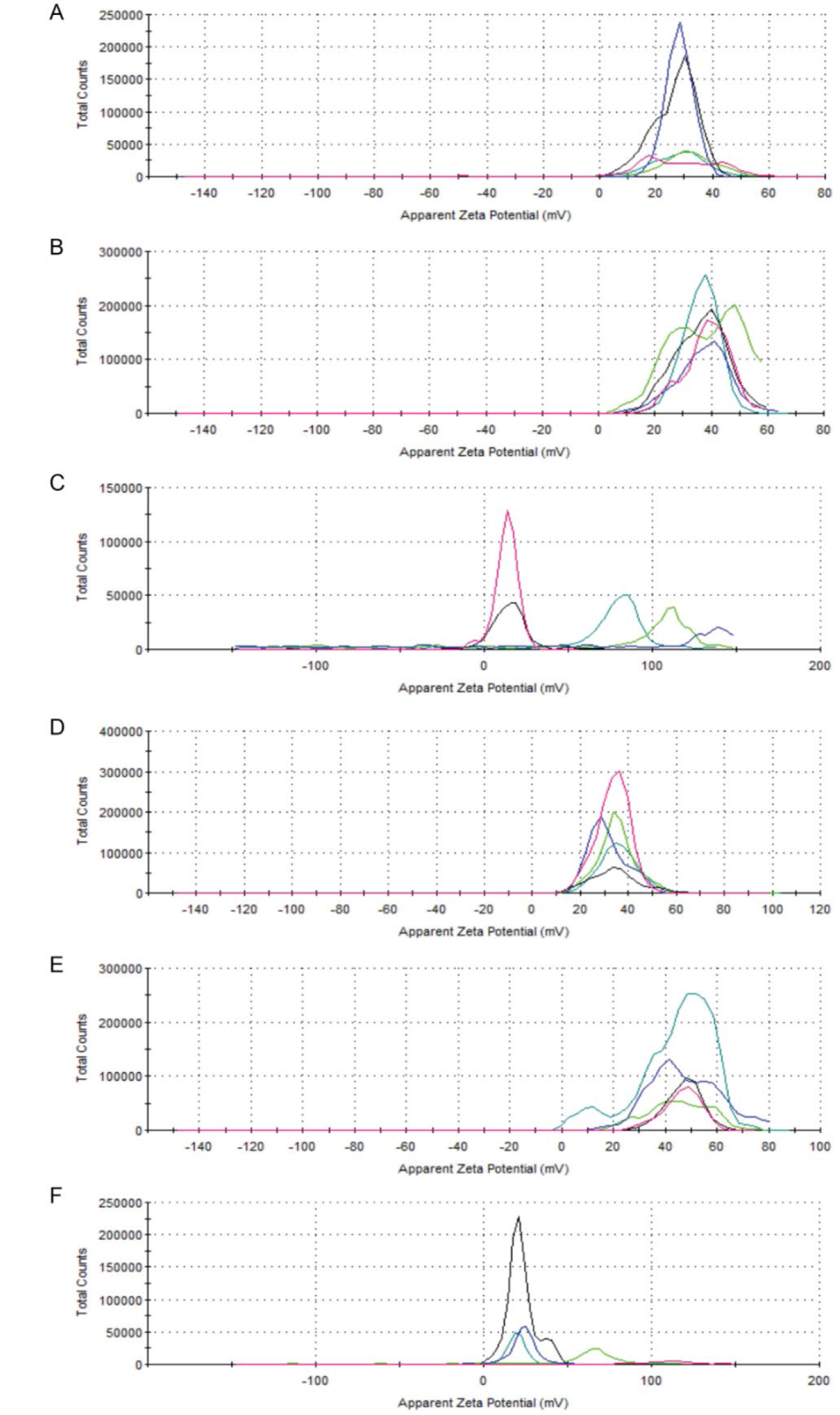
Particle charge distribution for nanoparticles self-assembled by peptides 1-6. **A.** particle charge distribution of nanoparticles self-assembled by peptide 1; **B.** particle charge distribution of nanoparticles self-assembled by peptide 2; **C.** particle charge distribution of nanoparticles self-assembled by peptide 3; **D.** particle charge distribution of nanoparticles self-assembled by peptide 4; **E.** particle charge distribution of nanoparticles self-assembled by peptide 5; **F.** particle charge distribution of nanoparticles self-assembled by peptide 6. Five independent replicated measurements per compound recorded by dynamic light scattering (DLS).

**Figure S7.**
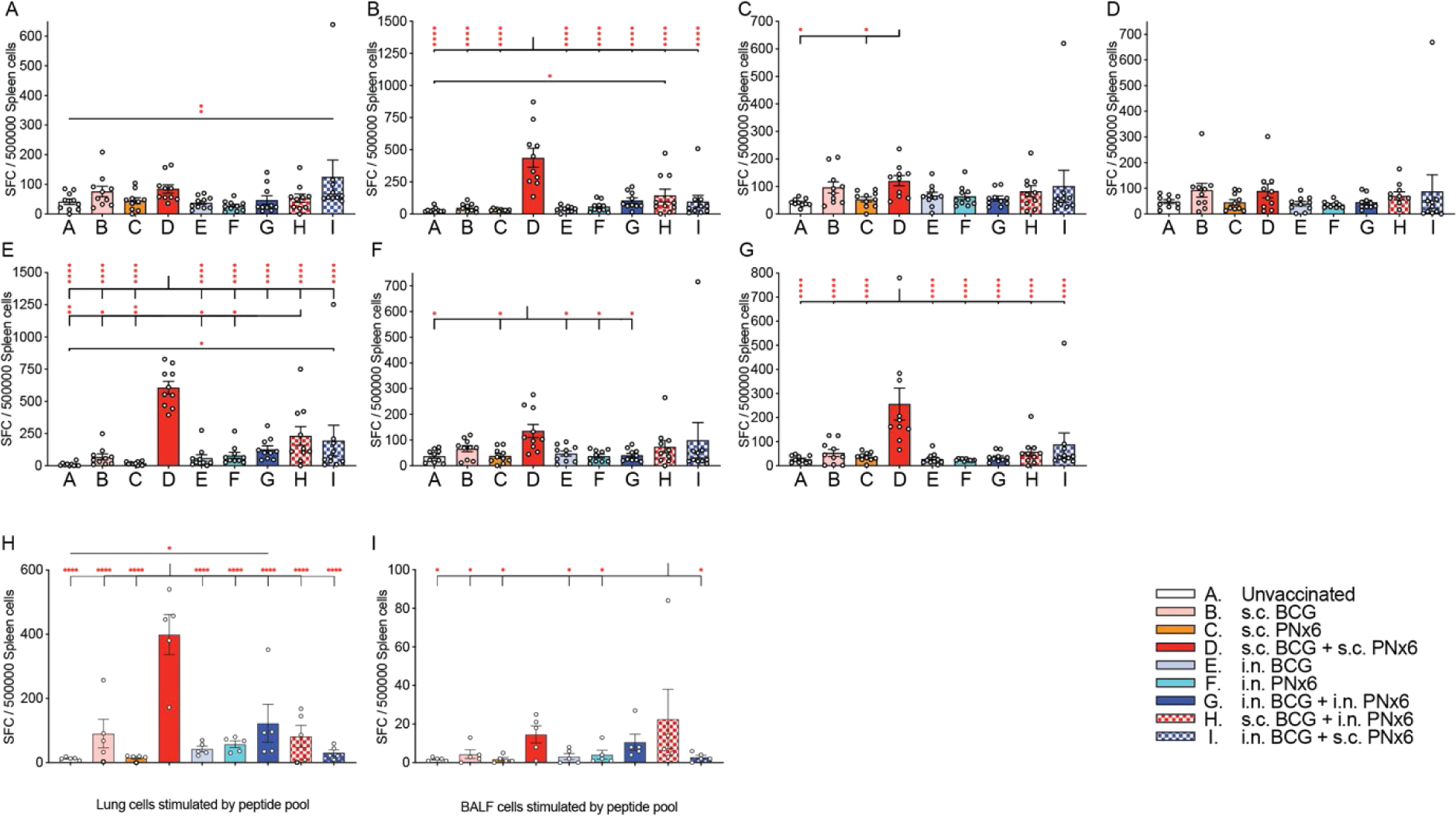
Secretion of IFNγ by spleen cells in response to stimulation with epitopes 1-6 and HLA-E binding peptide. **A.** SFCs of splenocytes stimulated by epitope 1; **B.** SFCs of splenocytes stimulated by epitope 2; **C.** SFCs of splenocytes stimulated by epitope 3; **D.** SFCs of splenocytes stimulated by epitope 4; **E.** SFCs of splenocytes stimulated by epitope 5; **F.** SFCs of splenocytes stimulated by epitope 6; **G.** SFCs of splenocytes stimulated by HLA-E binding peptide; **H.** SFCs of lung cells stimulated by epitope 1-6 mixture; **I.** SFCs of BALF cells stimulated by epitope 1-6 mixture. Each circle in **A**-**I** represents an individual mouse. Differences between groups in **A**-**G** were tested using one-way ANOVA with Fisher’s LSD multiple comparison. All data show means ± SEM or individual mice. *P< 0.05; **P< 0.002; ***P< 0.0002; ****P< 0.0001.

**Figure S8.**
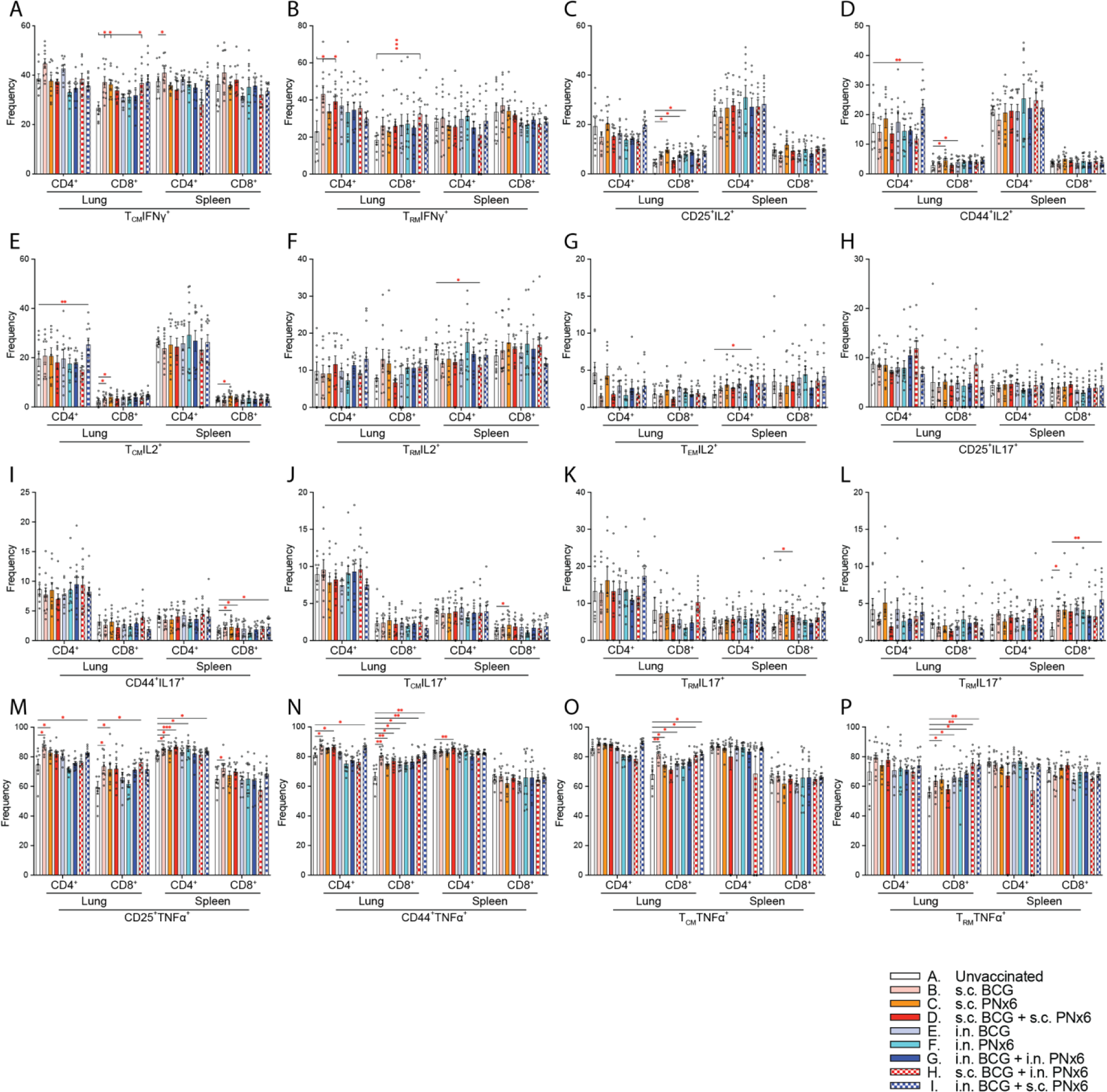
Frequency of different T cell phenotypes producing IFNγ, IL2, IL17 and TNFα in lung and spleen. **A.** Frequency of IFNγ producing CD4^+^ and CD8^+^ lung and spleen T_CM_ cells; **B.** Frequency of IFNγ producing CD4^+^ and CD8^+^ lung and spleen T_RM_ cells; **C.** Frequency of IL2 producing CD4^+^ and CD8^+^ lung and spleen CD25^+^ T cells; **D.** Frequency of IL2 producing CD4^+^ and CD8^+^ lung and spleen memory T cells; **E.** Frequency of IL2 producing CD4^+^ and CD8^+^ lung and spleen T_CM_ cells; **F.** Frequency of IL2 producing CD4^+^ and CD8^+^ lung and spleen T_RM_ cells; **G.** Frequency of IL2 producing CD4^+^ and CD8^+^ lung and spleen T_EM_ cells; **H.** Frequency of IL17 producing CD4^+^ and CD8^+^ lung and spleen CD25^+^T cells; **I.** Frequency of IL17 producing CD4^+^ and CD8^+^ lung and spleen memory T cells; **J.** Frequency of IL17 producing CD4^+^ and CD8^+^ lung and spleen T_CM_ cells; **K.** Frequency of IL17 producing CD4^+^ and CD8^+^ lung and spleen T_RM_ cells; **L.** Frequency of IL17 producing CD4^+^ and CD8^+^ lung and spleen T_EM_ cells; **M.** Frequency of TNFα producing CD4^+^ and CD8^+^ lung and spleen CD25^+^ T cells; **N.** Frequency of TNFα producing CD4^+^ and CD8^+^ lung and spleen memory T cells; **O.** Frequency of TNFα producing CD4^+^ and CD8^+^ lung and spleen T_CM_ cells; **P.** Frequency of TNFα producing CD4^+^ and CD8^+^ lung and spleen T_RM_ cells. Each circle in **A**-**P** represents an individual mouse. Differences between groups in **A**-**P** were tested using the mixed-effects analysis with Fisher’s LSD multiple comparison. All data show means ± SEM or individual mice. *P< 0.05; **P< 0.002; ***P< 0.0002; ****P< 0.0001.

**Figure S9.**
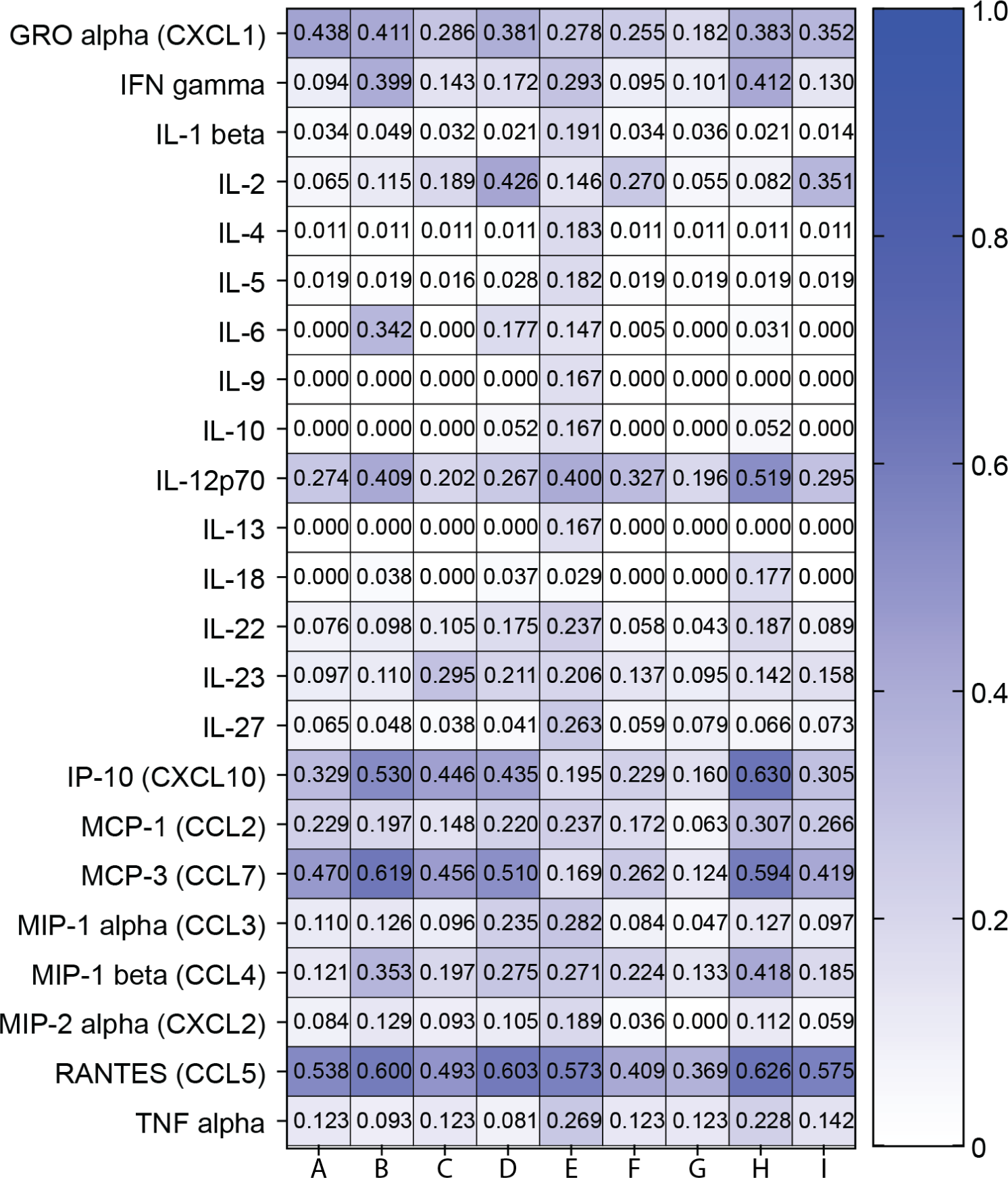
Profile of cytokines and chemokines in serum of vaccinated mice.

**Figure S10.**
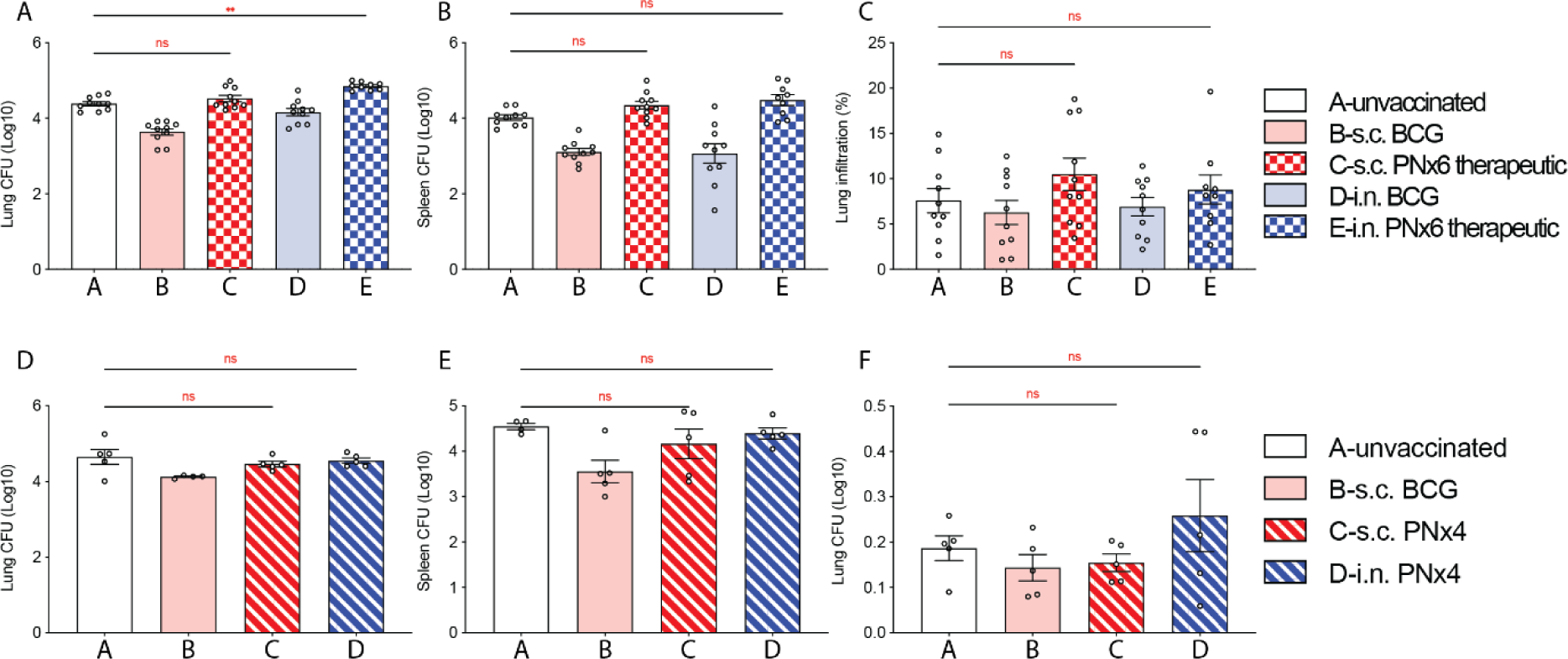
PNx4 and therapeutic use of PNx6 did not confer protection against TB. C57BL/6 mice were vaccinated with s.c. BCG, i.n. BCG s.c. PNx6 or i.n. PNx6. Sixty days later, mice were aerosol infected with 50-200 CFU of virulent Mtb H37Rv. 15 days later, mice were immunized with s.c. PNx6 or i.n. PNx6. Mtb bacterial burden was determined 45 days post infection in lungs and spleen. Lung damage was determined as the percentage of the area of dense cell infiltration in H&E-stained lung sections. **A.** Bacterial burden in lung; **B.** Bacterial burden in spleen; **C.** Cell infiltrates in lung. C57BL/6 mice were vaccinated with s.c. BCG, s.c. PNx4 or i.n. PNx4. Sixty days later, mice were aerosol infected with 50-200 CFU of virulent Mtb H37Rv. Mtb bacterial burden was determined 45 days post infection in lungs and spleen. Lung damage was determined as the percentage of the area of dense cell infiltration in H&E-stained lung sections. **D.** Bacterial burden in lung; **E.** Bacterial burden in spleen; **F.** Cell infiltrates in lung. Each circle in **A**-**F** represents an individual mouse. Differences between groups in **A**-**F** were tested using the one-way ANOVA analysis with Tukey’s multiple comparison. All data show means ± SEM or individual mice. *P< 0.05; **P< 0.002; ***P< 0.0002; ****P< 0.0001.

